# Identification of cambium stem cell factors and their positioning mechanism

**DOI:** 10.1101/2023.07.20.549889

**Authors:** Gugan Eswaran, Jacob Pieter Rutten, Jingyi Han, Hiroyuki Iida, Jennifer Lopez Ortiz, Riikka Mäkilä, Brecht Wybouw, Leo Vainio, Alexis Porcher, Marina Leal Gavarron, Jing Zhang, Tiina Blomster, Xin Wang, David Dolan, Ondřej Smetana, Siobhán M. Brady, Melis Kucukoglu Topcu, Kirsten ten Tusscher, J. Peter Etchells, Ari Pekka Mähönen

## Abstract

Wood constitutes the majority of terrestrial biomass. Composed of xylem, it arises from one side of the vascular cambium, a bifacial stem cell niche that also produces phloem on the opposing side. It is currently unknown which molecular factors endow cambium stem cell identity. Here we show that TDIF ligand-activated PXY receptors promote the expression of CAMBIUM AINTEGUMENTA-LIKE (CAIL) transcription factors to define cambium stem cell identity in the *Arabidopsis* root. By sequestrating the phloem-originated TDIF, xylem-dependent PXY confines the TDIF signaling front, resulting in the activation of CAIL expression and stem cell identity in only a narrow domain. Our findings show how signals emanating from cells on opposing sides ensure robust yet dynamically adjustable positioning of a bifacial stem cell.

## Main Text

In seed plants, stem cell populations that drive apical-basal growth are formed in the embryo. However, the vascular cambium, which promotes radial growth, and thus the majority of plant biomass, is defined *de novo* following germination (*1, 2*). Cambium is formed when cells with xylem identity promote stem cell function in their neighbors. These xylem identity cells are thus considered to be the cambium organizer cells (*3*) (**Fig. 1A**). The cambium is dynamic in size, ranging from a few to multiple undifferentiated cells concomitant with the level of active growth, yet contains only a single bifacial stem cell (*3*–*5*). How the organizer cells can exert their exquisite control over stem cells at variable distance is still unknown. A prerequisite to addressing this question is identification of regulators that define stem cell identity within the cambium, which have not been determined either. In the *Arabidopsis thaliana* root cambium, stem cell organizer is defined by a local signaling maximum of auxin, which contributes to stem cell positioning (*3, 6*). Auxin promotes the expression of CLASS III HOMEODOMAIN-LEUCINE ZIPPER (HD-ZIP III) transcription factors defining xylem identity (*3, 7, 8*), as well as a receptor kinase, PHLOEM INTERCALATED WITH XYLEM (PXY) (*3*). TRACHEARY ELEMENT DIFFERENTIATION INHIBITORY FACTOR (TDIF), which is derived from phloem-expressed CLAVATA3/ESR-RELATED 41 (CLE41) and CLE44, is the cognate ligand of PXY. Disruption of TDIF-PXY signaling causes major patterning and stem cell maintenance defects (*9*–*14*). These defects suggest that the illusive regulators of cambium stem cell identity are likely to be TDIF-PXY regulated.

**Fig. 1.**
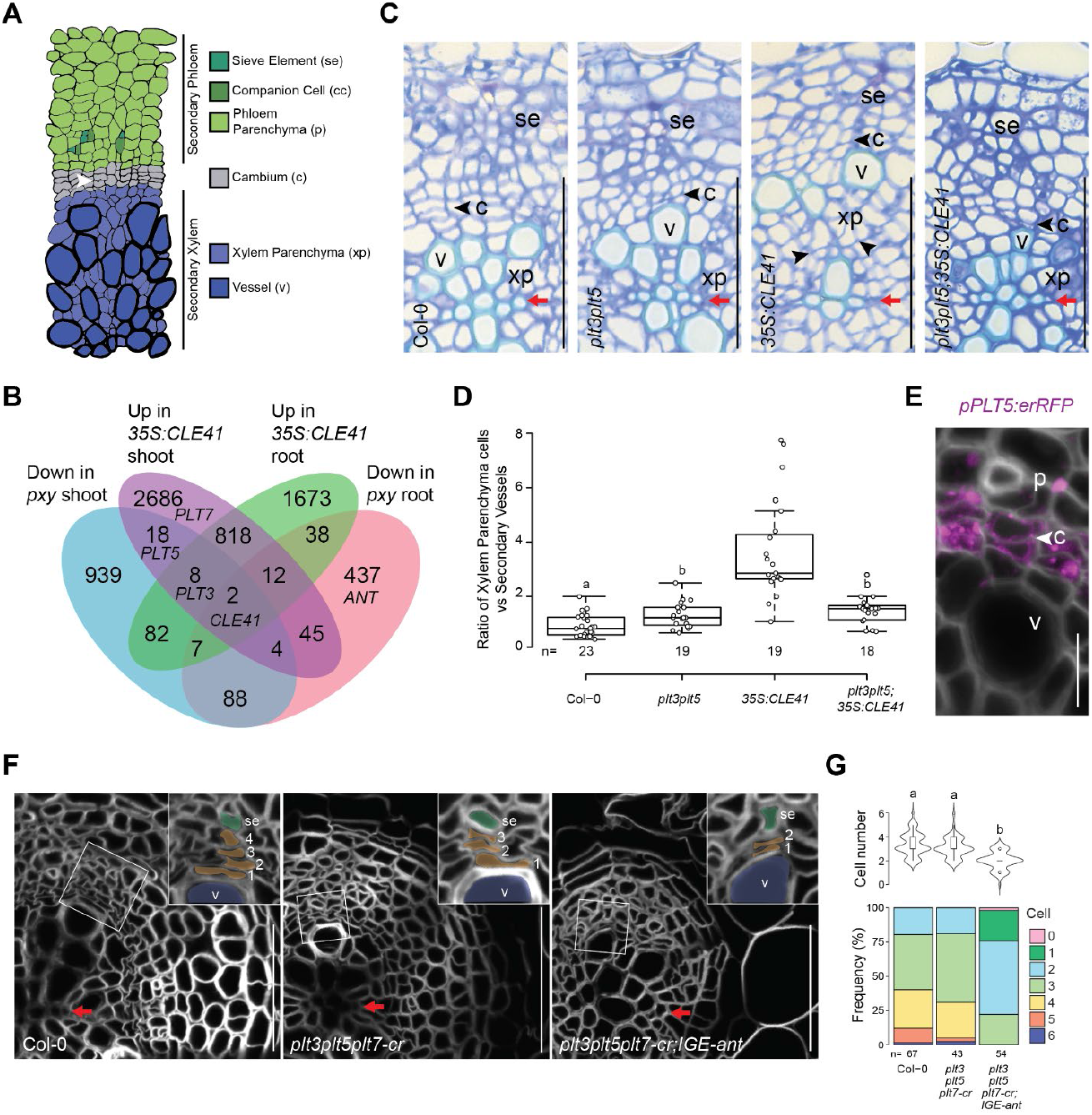
CAILs operate downstream of TDIF-PXY to maintain cambial stem cells. **(A)**Schematic representation of 28-day-old wild-type (Col-0) cross section of *Arabidopsis* root. **(B)**Venn diagram showing numbers of genes with differential expression in *pxy* and *35S:CLE41* shoots or roots relative to Col-0 in 7-day-old seedlings. Genes upregulated in *35S:CLE41* and down-regulated in *pxy*, were considered more likely to be PXY-signaling targets. Sections within the Venn diagram where *PLT3, PLT5, PLT7*, and *ANT* fall are marked. **(C)** Cross-sections of vascular tissue root morphology in Col-0, *plt3plt5, 35S:CLE41*, and *plt3plt5;35S:CLE41* (14-day-old seedlings). **(D)** Ratio between numbers of xylem parenchyma cells and secondary vessels. **(E)** Confocal cross-section of 14-day-old *pPLT5:erRFP* root. **(F)** Confocal root cross-sections of Col-0, *plt3plt5plt7-cr* and *plt3plt5plt7-cr;IGE-ant*. The false coloring in the inset highlights the cell types used for the quantification in panel (G). **(G)** Upper panel shows violin plot of cell number distribution between differentiated vessels and sieve elements, as visualized in the insets of panel (F). Lower panel shows the same data as distribution of cell number frequency between differentiated vessels and sieve elements. Letters in (D and G) indicate significant differences using Kruskal-Wallis test followed by Dunn’s post-hoc multiple comparisons with Bonferroni adjusted *p-values* (*p* < 0.05). Black/White arrowheads mark recent cell division, red arrows mark the primary xylem axis. Phloem (p), sieve element (se), cambium (c), xylem vessels (v), and xylem parenchyma (xp). Scale bars, 50 μm (C, F) and 10 μm (E).

### Identification of CAILs as cambium stem cell factors downstream of TDIF-PXY

To identify genes involved in specifying cambium stem cells, we compared transcriptomes of wild type *Arabidopsis* seedlings undergoing cambium initiation to an overexpressor of TDIF (*35S:CLE41*) and *pxy* mutant, characterized by enhanced or reduced TDIF-PXY signaling, respectively (*9, 15*). In line with these phenotypes, GO terms under-represented in *pxy* included meristem initiation; those over-represented in *35S:CLE41* included meristem growth (**fig. S1**). Transcripts under-represented in *pxy* and over-represented in *35S:CLE41* included *PLETHORA 3* and *5 (PLT3* and *PLT5*), members of the *AINTEGUMENTA-LIKE/PLT* (*AIL/PLT*) family (**Fig. 1B**). Previously, different members of the AIL/PLT transcription factor family have been associated with promoting stemness and/or an undifferentiated state in apical meristems (*16*–*19*). In cambium, *ANT* is specifically expressed in stem cells (*3*) and its absence leads to reduced radial growth (*20*). Thus, AIL/PLTs represent strong candidates for cambial stem cell regulators. Ectopic xylem parenchyma cell proliferation phenotype of *35S:CLE41* was suppressed by *plt3plt5*, demonstrating that these genes are required for cambium stem cell over-proliferation in *35S:CLE41* (**Fig. 1, C and D**). Nevertheless, the *plt3plt5* line demonstrated no obvious cambium phenotype (**Fig. 1, C and D**). Thus, we investigated the AIL/PLT family for further redundancy. Expression analysis of *AIL*/*PLT* fluorescent reporters showed that *ANT* (*3*), *PLT3, PLT5* and *PLT7* are expressed in cambium, while *PLT1, PLT2* and *PLT4* appear to be absent (**Fig. 1E; and fig. S2**). *ANT* was downregulated in the *pxy* mutant and *PLT7* upregulated in *35S:CLE41* (**Fig. 1B**). These data together suggest that these four CAMBIUM AILs (CAILs), PLT3, PLT5, PLT7 and ANT, operate redundantly in cambium development downstream of TDIF-PXY signaling. While *ant* mutants were characterized by reductions in cambium activity (*20*), *plt3plt5* double and *plt7* single mutants failed to show cambial phenotypes (**Fig. 1, C and D; and fig. S3A**). By contrast, a *plt3plt5plt7* triple null mutant (*plt3plt5plt7*-cr) (*21*) had reduced radial growth (**fig. S3B**). To determine levels of redundancy between *ant* and *plt*s, several mutant combinations were analyzed. *antplt3plt7* roots demonstrated occasional differentiation of cambial cells (**fig. S3C**). However, this line is unable to maintain the shoot apical meristem (*16*) and quadruple CAIL mutants did not survive to form cambium (**Materials and Methods**). Thus, to both assess loss of the four CAIL genes during cambium development, and to avoid secondary effects caused by shoot apical meristem loss, we generated an inducible genome editing (IGE) construct (*22*) targeting *ANT* (*ANT-IGE*). *ANT-IGE* was introduced to both the null *plt3plt5plt7-cr*(*21*) and *plt3plt5plt7-tdna* (*23*) backgrounds (**Fig. 1F; and fig. S3D)**. Root and shoot morphology appeared normal in both conditional quadruple mutants, albeit *plt3plt5plt7-cr;IGE-ant* showed slightly reduced root length (**fig. S3E**). *plt3plt5plt7-tdna;IGE-ant* displayed radial sectors without differentiated secondary xylem vessels and reduced phloem sieve elements suggesting loss of cambium identity (**fig. S3D**). In the *plt3plt5plt7-cr,IGE-ant* null background, secondary growth was significantly reduced (**fig. S3B**), and this was associated with a reduction in cambial cells per cell file, or occasionally, cell files with a complete loss of cambial cells (**Fig. 1, F and G**). These data demonstrate that the four CAILs are critical in maintaining vascular cambium identity.

The expression of *CAIL*s is specific to the cambial stem cells and their neighboring daughters. In these daughter cells an overlap between *CAIL*s and early xylem and phloem identity genes occurs (*3*) (**Fig. 2A; and fig. S4A**). To investigate the role of CAILs in shaping these boundaries, we focused on PLT5 as a representative factor. To obtain a genome-wide view of PLT5 action in the cambium, we performed RNA-seq on root tissues undergoing radial growth after 8 hours and 24 hours of induced overexpression of PLT5. We generated a 17-β-estradiol inducible (*24*) line, *35S:XVE>>PLT5-TagRFP*, for that purpose. Remarkably, 37% of xylem identity (*25*) and 53% phloem identity (*26*) genes were downregulated after 8 hours of *PLT5* induction, in comparison to 26% of all genes downregulated. Among core cell cycle genes (*27*) 55% were upregulated after 24 hours of PLT5 overexpression in comparison to 36% of all upregulated genes (**Fig. 2B; fig. S5, A and B**). Thus, PLT5 regulates large set of genes associated with cell proliferation, and xylem and phloem formation, upregulating the former while downregulating the latter two.

**Fig. 2.**
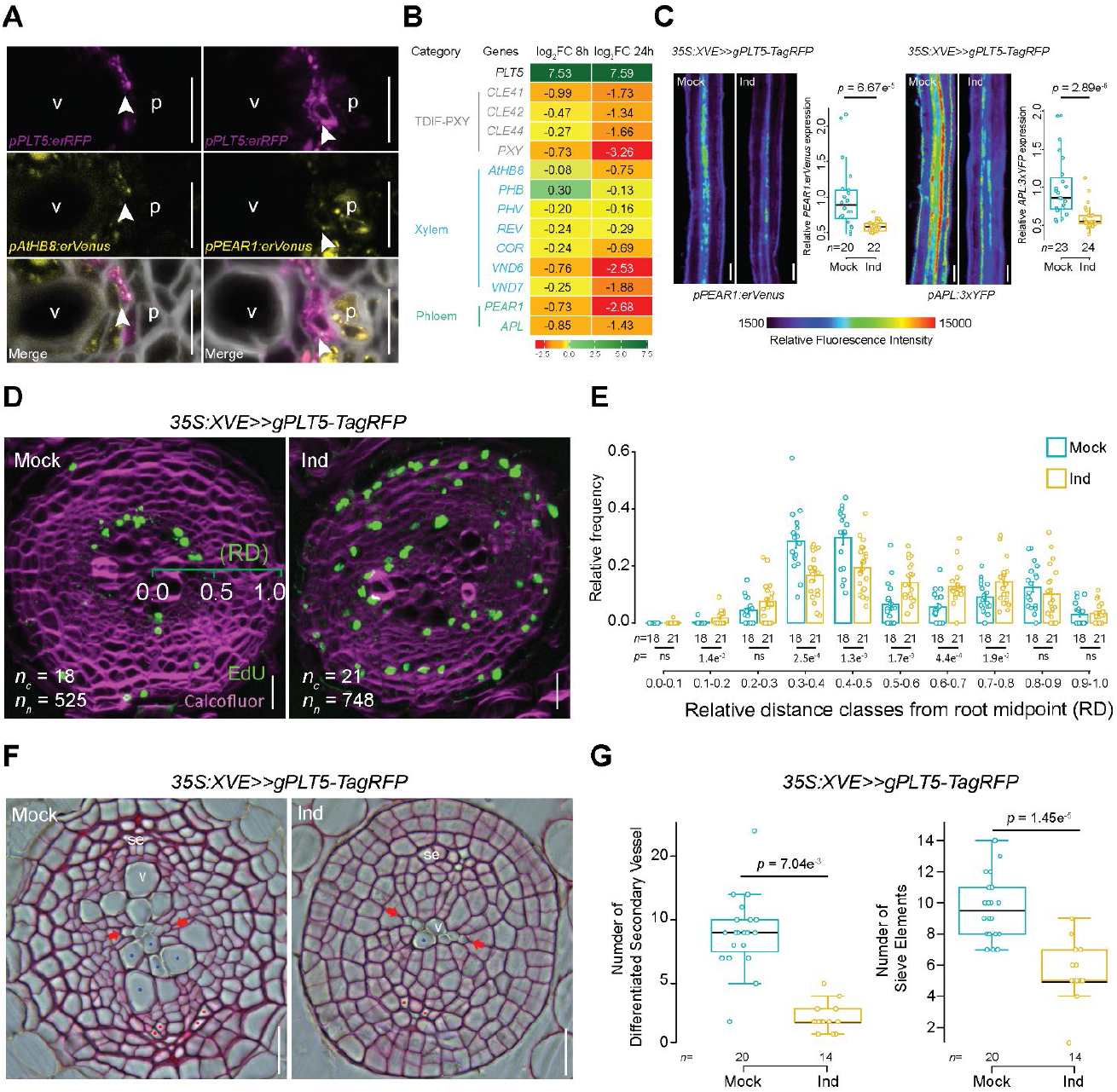
PLT5 promote cambial cell divisions and inhibit secondary xylem and phloem differentiation. **(A)** Confocal cross-sections of 12-day-old *pPLT5:erRFP pATHB8:erVenus* (left) and *pPLT5:erRFP pPEAR1:erVenus* (right) double marker lines. **(B)** Heatmap showing normalized log2FoldChange (FC) as determined by RNA-seq, upon induction of *PLT5* over-expression for 8 hours and 24 hours. **(C)** Lateral view of 8-day-old *35S:XVE>>gPLT5-TagRFP* roots with early (*pPEAR1:erVenus*) and late (*pAPL:3xYFP*) phloem markers after a 2-day induction. Box plots show quantification of relative fluorescent signal intensity. **(D)** Confocal cross-sections of *35S:XVE>>gPLT5-TagRFP* after 2-day induction with 16 hours of EdU (green) incorporation (in 12-day-old roots) to detect S-phase nuclei. **(E)** Barplot of relative distribution (RD) of EdU-positive nuclei along the radial root axis in panel (D) (primary xylem axis = 0; root surface = 1). Scale shown on green on the mock panel; *n*_c_ = number of cross-sections analyzed; *n*_n_ = total number of nuclei. **(F)** Cross-sections of *35S:XVE>>gPLT5-TagRFP* after 4-day induction (in 8-day-old plants). Red dots indicate the sieve elements and blue dots indicate vessels. **(G)** Quantification of secondary xylem vessel and sieve element numbers. In (E) error bars show mean ± standard error. Significant differences were calculated using two-tailed student t-test for each relative distance class (ns = *p*>0.05). In (C) and (G) significant differences between mock and induced conditions were calculated using two-sided Wilcoxon-Mann-Whitney test. White arrowheads mark recent cell division, red arrows mark the primary xylem axis. Vessels (v), phloem (p), sieve elements (se). Scale bars, 10 μm (A), 100 μm (C), and 50 μm (D and F).

Next, we investigated the consequences of *PLT5* induction on morphogenesis. Short-term *PLT5* induction promoted ectopic DNA replication observed with EdU staining, accompanied by ectopic cell divisions within xylem and phloem (**Fig. 2, D and E; and fig. S5C**). In support of our RNA-seq data (**Fig. 2B; fig. S5, A and B**), *PLT5* induction caused rapid down-regulation of xylem (*VND6*) and phloem (*PEAR1, APL*) fluorescent reporter lines (**fig. S5D; and Fig. 2C**), leading to inhibition of xylem vessel and phloem sieve element formation (**Fig. 2, F and G**), and subsequently, inhibition of radial growth (**fig. S5E**). These data demonstrate that in the cambium, PLT5 maintains cell division capacity, and the undifferentiated state of cambium cells. This occurs both through promoting cell division and through active opposition of differentiation to either of xylem or phloem fate. Together, over-expression and loss-of-function analysis shows that CAILs are key cambium stem cell factors.

### Computational model for cell fate determination in cambium

*CAIL* expression occurs only in a narrow stem cell domain of the cambium. This is in contrast to the *PXY* receptor expression domain, which is strongest in the xylem/organizer domain tapering off towards weak expression in stem cells (*3*) **(Fig. 3A; and fig. S4B**). This raised the question of why the downstream CAILs are constrained to the subdomain of low *PXY* expression. In any biological system, ligand binding to a receptor results in its sequestration from the pool of free ligands. *CLE41* expression and the subsequent TDIF peptide gradient extends from the phloem (*13, 15, 28*) (**fig. S4C**), thus the first PXY receptors TDIF peptides encounter are those located at the lower end of the PXY gradient. Therefore, we hypothesized that sufficiently strong TDIF sequestration by PXY could abrogate further TDIF spread and thereby lead to a narrow active TDIF-PXY signaling domain and hence restrict CAIL expression to the stem cells. After 24 hours application of synthetic TDIF peptide, the expression of *PLT5* expanded towards xylem parenchyma, coinciding with ectopic cell divisions (**Fig. 3A**), supporting the idea that excess TDIF prohibits sufficient sequestration. Long-term TDIF treatment led to further expansion of *PLT5* expression and cell proliferation in xylem parenchyma (**Fig. 3B**).

**Fig. 3.**
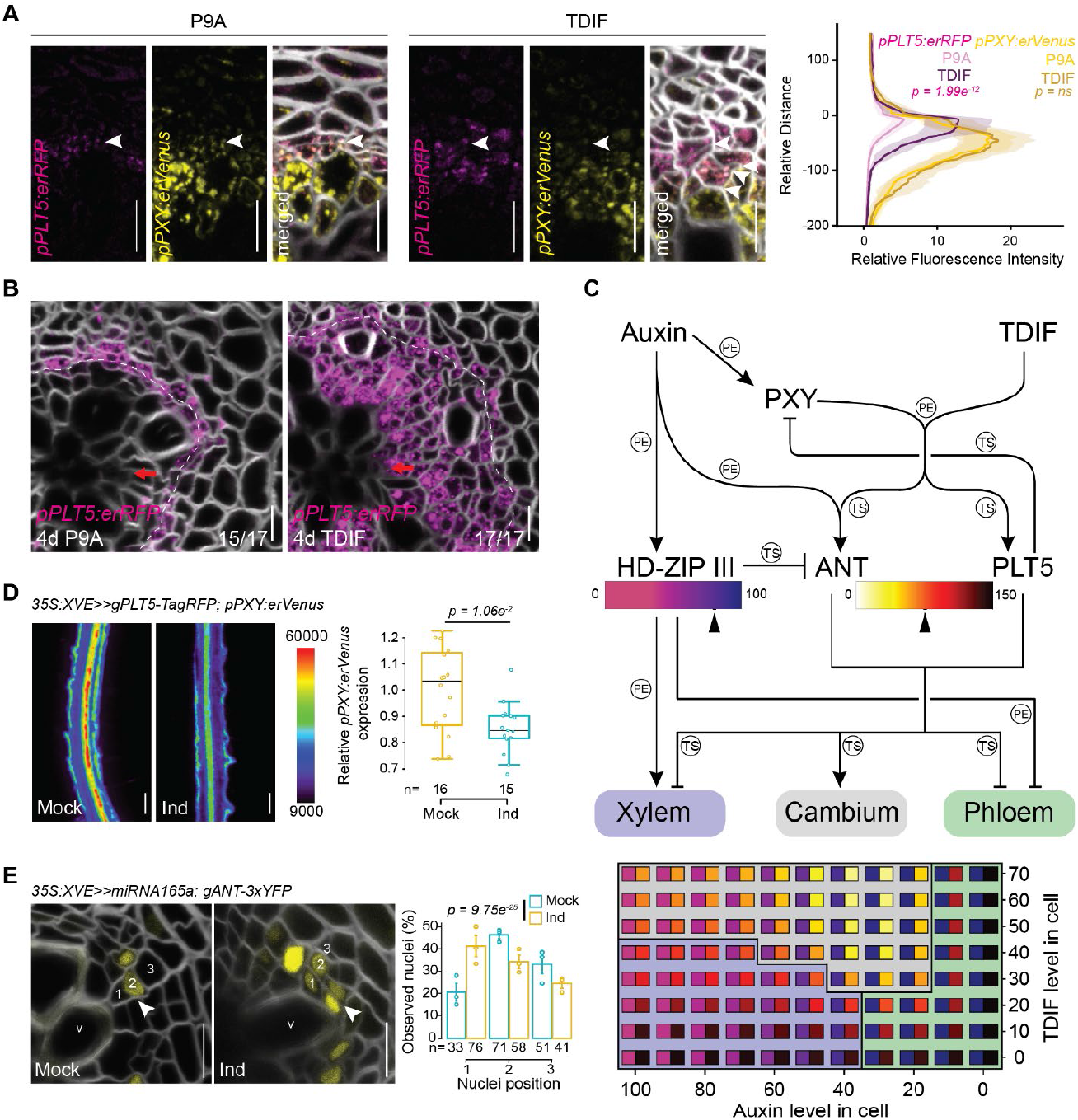
Regulatory network defining cambium, xylem and phloem. **(A)** Confocal cross-sections of roots carrying *pPLT5:erRFP* and *pPXY:erVenus* double markers in 15-day-old plants after 24 hours of P9A_(n=18)_ (control) or TDIF_(n=16)_ treatment. P9A is a mutant version of TDIF, incapable to bind PXY. While wild-type or P9A-treated cambia typically have a single cell in every radial cell file having undergone a recent division (thin cell wall, marked with white arrowhead), TDIF-treated cambium have additional, ectopic divisions in xylem domain (multiple arrowheads). Quantification of *pPLT5:erRFP* expression (right) shows increases and a shift towards xylem. (ns = *p*>0.05) **(B)** Confocal cross-sections of *pPLT5:erRFP* expression after 4-day TDIF treatment (in 13-day-old root). White dashed line marks the cambium. **(C)** Modelling cambial cell fate decision making under different combinations of auxin and TDIF concentrations. Top panel: Interaction network underlying cambial cell fate decision making incorporated in the model. PE, refers to ‘<u>P</u>ublished <u>E</u>lsewhere’, TS refers to ‘<u>T</u>his <u>S</u>tudy’. Supporting details with references are listed in (fig. S6). Bottom panel: Fate map of a cambial cell exposed to different auxin-TDIF combinations, for each Auxin-TDIF combination, the left square within the rectangle shows the summed PLT+ANT level at steady state and the right square shows the HD-ZIP III level at steady state. The black arrowheads on the color bars in the top panel show the threshold levels. Applied cell fate thresholds are as follows: If PLT+ANT >= 75 cambial identity (grey); if PLT+ANT < 75 and HD-ZIP III >= 30 xylem identity (blue); if HD-ZIP III < 30 phloem identity (green). Parameter settings are described in the Modelling Methods (strong HD-ZIP III repression settings). **(D)** Lateral view of *35S:XVE>>gPLT5-TagRFP;pPXY:erVenus* 8-day-old plants after a 2-day induction with quantification of relative fluorescent signal intensity. Significant difference between mock and induced condition was calculated using two-sided Wilcoxon-Mann-Whitney test. **(E)** Confocal cross-sections of *35S:XVE>>miRNA165a;gANT-3xYFP* after 2-day induction (in 12-day-old plants). Quantification of positional expression of *gANT-3xYFP* in cambium. Barplot showing position of *gANT-3xYFP*-positive nuclei. In (E) categorical distribution was tested using a chi-square test. Data points represent the mean of each biological repeat. White arrowheads mark recent cell division, red arrows mark the primary xylem axis. Vessels (v). Scale bars, 10 μm (A, B and E), 200 μm (D).

To address whether the TDIF sequestration hypothesis could explain the above observations, we developed a computational model. This model combines the regulatory interactions described here and those published previously (**Fig. 3C, top; and fig. S6)**, to determine cambial cell division and differentiation dynamics **(Modelling Methods**). Previous models investigated 2D patterning dynamics while partly invoking hypothetical regulatory factors (*29*–*31*). Instead, we focused on filling in missing links explaining cell fate decisions by using a simple 1D static tissue model. The model incorporates TDIF-PXY promotion of *PLT5* (as a representative of the PLT subclade) and *ANT* (**Fig. 3A; and fig. S4B**); repression of *PXY* by PLT5 which we observed upon induction of *PLT5* (**Fig. 2B; and 3D**); auxin-mediated promotion of HD-ZIP III (*3, 7, 8*), ANT transcription factor (*3, 32*), and PXY receptor (*3*); and HD-ZIP III promotion of xylem differentiation (*3, 33*– *35*) and inhibition of phloem differentiation (*3, 36*). By using *HD-ZIP III*-targeting inducible miR165a (*3*), we noticed that HD-ZIP IIIs repress *ANT* in xylem identity cells (**Fig. 3E**). Prior to testing the influence of TDIF sequestration by PXY in a multicellular setting, we first explored the capacity of this network to correctly assign cell fate identity in a single cell given various auxin and TDIF levels. Specifically, (i) cells experiencing high auxin and low TDIF levels should express HD-ZIP III and acquire xylem fate; (ii) cells with high TDIF and low auxin levels should obtain phloem fate; and (iii) cells with intermediate TDIF and auxin levels should have high ANT and PLT expression and thus cambium identity. Parameter sweeps showed that these requirements were met for a wide range of parameter values independently of specific fate determination threshold levels (**fig. S7, A and B; and Modelling Methods)**. However, in the presence of high TDIF and high auxin levels, a more robust formation of the xylem occurs in conditions when ANT is strongly repressed by HD-ZIP III (**fig. S7C; and Fig. 3C, bottom panel**).

### TDIF sequestration mechanism explains observed cambium phenotypes

Next, we extended our model to a row of 3-5 cells to represent a cambium cell file. An auxin gradient with a maximum in the xylem identity cell and a TDIF gradient with a maximum in the phloem identity cell were superimposed. For strong HD-ZIP III repression of *ANT*, correct patterning of the three cell types occurred readily under variable TDIF gradient settings (**fig. S8A**). However, when varying cell number, in a 3-cell system, intermediate TDIF production resulted in invasion of PLT expression into the xylem, while the same settings for 5 cells resulted in a TDIF-PXY overlap insufficient for promoting PLT5 expression and subsequent cambium identity (**fig. S8B**). Clearly, increasing or decreasing of TDIF production would improve one issue at the cost of the other, implying that strong HD-ZIP III-mediated ANT repression results in insufficient patterning robustness. We thus turned to our hypothesis described above, testing for an important role of strong sequestration of TDIF upon binding to PXY (**fig. S8C**). Here, free movement of TDIF occurs until it encounters sufficient PXY receptors to effectively halt its diffusion. While this resolved the TDIF-PXY dependent invasion of PLT5 expression into the xylem for 3 cells, absence of strong HD-ZIP III repression of the partly auxin induced ANT also caused an invasion of ANT into the xylem (**fig. S8C**). Combined, this suggests that both mechanisms must be active *in planta* to some extent. To test which of these mechanisms dominates patterning of a narrow cambial domain observed *in planta*, we created parameter conditions under which they could be tested in isolation. Since both mechanisms create a narrow *ANT* and *PLT* expression domain under variable cell numbers in normal conditions (**Fig. 4A, leftmost column; and fig. S8**), we sought to investigate both hypotheses in perturbed conditions. One prominent difference between the modelled “strong HD-ZIP III”, and “TDIF sequestration” parameter regimes was the predicted behavior of ANT. In strong HD-ZIP III parameter regimes, the model predicted that lower *PXY* expression would reduce active TDIF-PXY complexes and consequently CAIL expression in the cambium (**Fig. 4A**). By contrast, in regimes where TDIF sequestration was dominant, the model predicted that reduced *PXY* expression resulted in xylemward expansion of TDIF gradient, leading to activation of further PXY receptors and hence ANT expression towards the xylem (**Fig. 4B**). To compare these differential predictions against *in planta* behavior, we reduced *PXY* by inducible RNA interference (*RNAi-PXY*) in *ANT* fluorescent reporter background in Arabidopsis roots. Upon *RNAi-PXY* induction, *PXY* levels dropped to 71% of wild type levels (**fig. S5F**) and a shift in *ANT* expression towards the xylem was observed (**Fig. 4, C and D**), as the model predicted for the TDIF sequestration scenario.

**Fig. 4.**
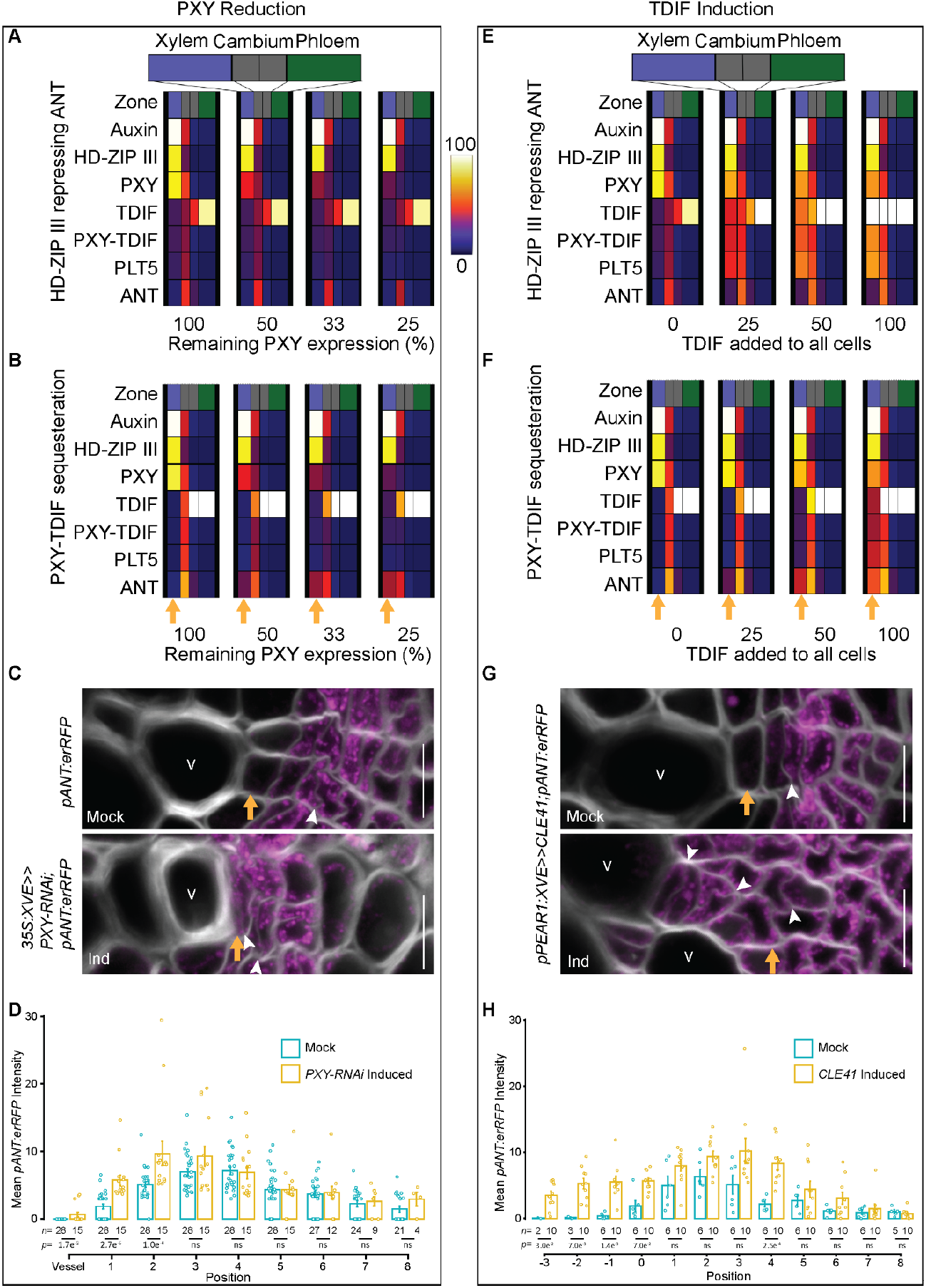
TDIF sequestration ensures narrow *ANT* expression domain. **(A, B, E** and **F)** Comparison between strong HD-ZIP III repression parameter regime (A and E) and strong sequestering effect of PXY on TDIF in a variable spatial context (B and F). Heatmap shows the gene expression level. Parameter settings are described in the Modelling Methods (strong HD-ZIP III repression versus strong sequestration settings). (A and B) Effect of reducing *PXY* expression in the two regimes. In (A) this leads to overall reduction of *ANT* expression, while in (B) it leads to a xylemward shift of *ANT* expression. (E and F) Responses of the two regimes to various levels of enhanced TDIF induction in phloem. Strong HD-ZIP III repression safeguards the xylem from ANT expression, while in the strong sequestering regime *ANT* is expressed in the xylem. **(C)** Confocal cross-sections of *35S:XVE>>PXY-RNAi; pANT:erRFP* after 2-day induction (in 12-day-old plants). **(D)** Quantification of mean *pANT:erRFP* intensity position. **(G)** Confocal cross-sections *pPEAR1:XVE>>CLE41;pANT:erRFP* after 2-day induction (in 12-day-old plants). **(H)** Quantification of mean *pANT:erRFP* intensity position. In (D and H) error bars show ± standard error. Significance was determined using two-sided Wilcoxon-Mann-Whitney test, (ns = *p*>0.05). White arrowheads mark recent cell divisions, orange arrows mark the position of xylem identity cells before induction. Vessels (v). Scale bars 10 μm (C and G).

Further corroboration of the dominance of TDIF sequestration came from the predicted differential responses of the two modelled mechanisms to increased TDIF levels emanating from the phloem. In the strong HD-ZIP III mechanism, a modelled increase in TDIF levels resulted in little change to the *ANT* expression domain (**Fig. 4E**). By contrast, for TDIF sequestration, expansion of the *ANT* expression domain towards the xylem was predicted (**Fig. 4F**). Supporting the sequestration model, TDIF overproduction *in planta* via phloem precursor-specific *pPEAR1:XVE>>CLE41* line resulted in rapid expansion of *ANT* expression into the xylem (**Fig. 4, G and H**). Thus, while both mechanisms may co-exist *in planta*, our modelling and experimentation suggest that the sequestration mechanism provides the dominant patterning constraint. Thus, the sequestration of TDIF by its PXY receptor effectively constrains TDIF mobility to the first few PXY expressing cells, enabling robust, spatially constrained patterning of the vascular cambium. The *in planta* mechanism underlying this strong sequestration remains to be elucidated but may arise either from sustained TDIF-PXY binding or alternatively from TDIF-PXY internalization and subsequent TDIF degradation.

## Discussion

Our combined experimental and modelling approach shows how PXY-mediated TDIF sequestration generates a robust patterning mechanism. By manipulating the auxin and TDIF gradients, we observed that the balance between the auxin gradient and the TDIF gradient determines the localization of the cambial stem cells as well as cambium size. We propose that this patterning mechanism flexibly enables adjustment of both phloem to xylem ratio and overall growth. A dominance of auxin localizes the cambium stem cell phloemward, leaving room for xylem cells to differentiate (*6*) **(fig. S9**). A TDIF dominated condition localizes the cambium xylemward, allowing phloem cells to differentiate, although the underlying connection to phloem differentiation remains to be studied. A strong combination of the two gradients allows for a larger cambium that sustains a larger total cell production (**Fig. 4G; and fig. S9**). A WUSCHEL-related HOMEOBOX gene, WOX4, has been shown to act downstream of TDIF-PXY signaling (*37, 38*). It remains to be studied how WOX4 is integrated with the signaling network described here, particularly because *WOX4* has a broader expression domain than *CAIL*s in the cambium (this paper and (*3*)). Also, it remains to be determined through which mechanism TDIF-PXY promotes *CAIL* transcription. Previously, we discovered a stem cell organizer at the xylem side of the cambium, defined by high levels of auxin signaling (*3*). Here, CAIL transcription factors were identified as the key stem cell factors operating downstream of the auxin-regulated PXY receptor. We elucidated how through sequestering TDIF ligands on the edge of the PXY gradient, the auxin-promoted organizer can induce CAILs and perform cambial stem cell patterning at a distance. Our findings suggest that like animals, plants use opposing morphogen gradients fine-tuned by sequestration-based feedback mechanisms to control precise positioning and cell fate decisions.

## Acknowledgements

We thank Viola Willemsen, Kalika Prasad and Beth Krizek for providing us published material, and Jan Traas for providing the unpublished entry clone containing the *AIL6/PLT3* promoter; and Ykä Helariutta for providing feedback on the manuscript. Confocal imaging was performed with help and using equipment of the Light Microscopy Unit (LMU), University of Helsinki. Special thanks to Mikko Herpola and Filipa Alexandra Silva for technical help.

## Author contributions

K.t.T., J.P.E. and A.P.M. conceived the project; G.E., H.I., J.L.O., O.S., S.B., M.K.T, J.P.E. and A.P.M., designed the experiments; G.E., J.H., H.I., J.L.O., R.M., B.W., J.Z., M.K.T. and J.P.E. performed the experiments; J.P.R., K.t.T. designed and performed the computational modelling; L.V. created the circle-unwrapping projections; A.P. performed statistical analysis; X.W. generated genetic material; M.L.G. provided preliminary data; T.B. and D.D. assisted with RNASeq analysis; G.E., J.P.R, K.t.T., J.P.E. and A.P.M. wrote the paper with input from all authors.

## Data and materials availability

All lines involved in this study are available upon reasonable request from the corresponding authors. Gene accession numbers are as follows: *ANT*, AT4G37750; *APL*, AT1G79430; *AtHB8*, AT4G32880; *CLE41*, AT3G24770; *MIR165A*, AT1G01183; *PEAR1*, AT2G37590; *PLT1*, AT3G20840; *PLT2*, AT1G51190; *PLT3*, AT5G10510; *PLT4*, AT5G17430; *PLT5*, AT5G57390; *PLT7*, AT5G65510; *PXY /TDR*, AT5G61480; *VND6*, AT5G62380; *PEX4 /UBC21* AT5G25760. The data of transcriptomes of *pxy* and *35S:CLE41* is available on GEO (accession number <u>GSE119872</u>).

## Supplementary Materials

### The Supplementary pdf includes

Experimental Materials and Methods Modelling Methods

Tables S1 to S6 Figs. S1 to S9

References (39–51)

### Other Supplementary Materials for this manuscript includes the following

Data S1 (Primers, constructs, seeds)

