## Supplementary Material for "Identification of cambium stem cell factors and their positioning mechanism"

Gugan Eswaran *et al*.

Corresponding authors:
Kirsten ten Tusscher,
J. Peter Etchells,
Ari Pekka Mähönen,

**The PDF file includes:**

Materials and Methods
Modelling Methods
Figs. S1 to S9
tables S1 to S6
References

**Other Supplementary Materials for this manuscript include the following:**

Data S1

### Materials and Methods

Plant material and cloning

The Col-0 background was used throughout. *pxy* (*9*), *35S:CLE41* (*12*)*, ant-GK* (GK-874H08) (*20*), *plt3plt5plt7-cr* (*21*), *plt3plt5plt7-tdna* (*23*), have been described previously.

All entry clones were generated by PCR amplification of the target sequence. PCR products were then recombined into MultiSite Gateway compatible pDONR entry vectors using a BP clonase reaction. Multisite Gateway technology was used to assemble the entry clones into Gateway compatible binary vectors using multisite gateway LR reactions. Primers for PCR amplification, pDONR entry vectors, and expression vectors are listed in the data S1.

Upon plant transformation of expression vectors, single insertion lines were identified based on Mendelian segregation of the selectable marker. Multiple single insertion lines were screened for each construct to observe the most consistent expression patterns or phenotypes.

In an attempt to generate higher order *plt/ant* mutants, we crossed *plt3plt5plt7-tdna* (♀) (*plt3-1*, SALK_127417, *plt5-2*, SALK_059254; *plt7-1*, SAIL_1167_C10) (*23*) x *antplt3plt7* (♂) (*ant-GK*, GK-874H08). Of 78 F1 seeds tested, none of them germinated, even when 11 of the seeds were attempted to germinate on $GA_3$. To generate the quadruple mutant with different alleles we crossed *plt3plt7* (♀) (*plt3-1*, SALK_127417, *plt7-1*, SAIL_1167_C10)(*23*) x *antplt5* (♂) (*ant-GK*, GK-874H08; *cho1-3)* (*39*)*,* resulting F1 plants heterozygous for all the four alleles. However, no homozygous plants for all the alleles could be identified in F2. With these alleles, in the segregating population we were unable to identify any higher order than double homozygous mutant combinations, due to likely gametophyte or embryo lethality. However, triple mutant combination of *plt3, plt5* and *plt7* can produce seeds (*21*, *23*). To avoid the lethality of the quadruple mutant, we generated a conditional *plt*/*ail* quadruple mutant by taking an advantage of an inducible genome editing system (IGE) (*22*). An inducible CRISPR construct targeting *ANT* was transformed into two different *plt3plt5plt7* mutant backgrounds, a null combination *plt3plt5plt7-cr* (*21*) and a weaker allele combination *plt3plt5plt7-tdna* (*23*). This vector contained two sgRNAs designed to create a large deletion in *ANT* upon treatment with 17-β-estradiol. The primers used to generate two fusions of a small nuclear RNA promoter and sgRNA (*pAtU3-sgRNA*) are listed in the data S1. The resulting PCR products were cloned into a *p2PR3-Bsa I-ccdB-Bsa I* entry vector using Golden Gate and Gibson assembly cloning methods thus concatenating the two *pAtU3b-sgRNA1-ANT and pAtU3b-sgRNA2-ANT* fragments. The IGE binary vector was generated in a single MultiSite Gateway LR reaction by combining a 17-β-oestrodiol-inducible *WOODEN LEG* (*WOL*) promoter (*24*), *Cas9p*, *2R3z-2x-pAtU3b-sgRNA-ANT* and destination vector *pFRm43GW* (*22*) (destination vector). To facilitate screening of transformed seeds, we used seed-specific RFP fluorescence provided by pFRm43GW. The *plt3plt5plt7-cr;IGE-ant* and *plt3plt5plt7-tdna;IGE-ant* lines were germinated directly in 1/2 GM plates supplemented with 5 µM 17-β-estradiol and grown for 15 days and 13 days respectively alongside Col-0 and their respective *plt3plt5plt7-cr, plt3plt5plt7-tdna* controls.

Plant growth and chemical treatments

All the plants were grown vertically in a plate in a 23°C growth chamber with 8 hours dark & 16 hours light cycle. Seeds were surface sterilized using 70% ethanol with Tween-20 (µl/ml) solution for 5 min with vortexing, followed by five washes in sterile Milli-Q (MQ) water. The sterilized seeds were stratified for 2 days at 4°C in darkness before plating them on 1/2 germination medium (GM) containing 0.5x Murashige and Skoog (MS) media with vitamins (Duchefa), 0.8% plant agar, 1% sucrose and 0.5 g/l MES pH 5.8. Alternatively, 0.5x MS (pH 5.8), 1% sucrose and 1% agar was used. The age of the plants was measured from when the plates were vertically positioned in the growth cabinet.

Aqueous 10 mM stocks of P9A and TDIF peptides (GeneCust) were prepared stored at the -80°C. 10 mM stocks of EdU (Thermo Fisher), dissolved in dimethyl sulfoxide (DMSO) was prepared and stored at -20°C. 17-β-estradiol (EST) (Sigma), was prepared as a 20 mM stock solution in DMSO and stored at -20°C.

Short term (24 hours or less) TDIF and P9A treatments were performed in liquid 1/2GM containing the respective peptide with the working concentration of 10 µM. Longer treatments were performed on 1/2GM plates. 17-β-estradiol induced gene expression was achieved by transferring plants onto plates containing 5 µM 17-β-estradiol or an equal volume of DMSO as a mock treatment, except in **Fig. 2F and fig. S5E** and *35S:XVE>>PLT5-TagRFP* RNA-Seq, 1 µM 17-β-estradiol was used. For EdU incorporation, plants were placed in liquid 1/2GM containing 10 µM EdU for 16 hours prior to fixation.

#### RNA-Seq profiling and data analysis

Transcriptomes of *pxy* and *35S:CLE41* in comparison to wild type were determined from seedlings grown on vertical plates for 7 days. Upon harvesting, seedlings were separated into root and shoot samples by separating plants at the root-hypocotyl junction. RNA and library preparation in biological quadruplicate was performed as described (*40*). 50 bp single end reads were obtained on the Illumina HiSeq 4000 platform. Sequencing was performed by the QB3 Genomics Facility, University of California, Berkeley. Quality checking and trimming of the raw FASTQs was performed with Kraken (*41*) followed by alignment to TAIR10 with Tophat2 (*42*). An average of 3.4M reads were obtained per treatment. Gene counting was performed with HTSeq (*43*), and differential gene expression was determined with DESeq2 (*44*). Cut-offs for differential expression was an adjusted p value < 0.05. The data is available on GEO (accession number GSE119872). GO analysis was performed on genes differentially expressed in either *35S:CLE41* or *pxy* relative to wild type (Col-0).

For PLT5 transcriptome analysis, *35S:XVE>>PLT5-TagRFP* seeds were germinated on 1/2 GM plates for 9 days or 9 days 16 hours, and then transferred to 1 µM 17-β-estradiol or DMSO plates for 24 hours or 8 hours, respectively. For each sample, 1.5 cm of root segments below the root-hypocotyl junction were collected from 15 individuals. Visible lateral roots were removed. RNA isolation, library preparation and data analysis were done as previously described (*45*) except single-end reads (86bp) and were mapped to *Arabidopsis* reference genome (TAIR 10.39). Differential expression between the mock and inductions was analyzed using the edgeR package (*46*). Subsequently, Pvalue < 0.05 was applied to identify differentially expressed genes (DEGs).

#### Thin sections, GUS staining, and light microscopy

All the root samples were sectioned 5mm below the hypocotyl junction unless mentioned otherwise. Samples were fixed overnight in 1% glutaraldehyde, 4% formaldehyde in 0.05 M sodium phosphate buffer pH 7.2, followed by dehydration through an ethanol series, and embedding in plastic resin using either Historesin (Leica) or JB4 (Polysciences). 3, 5 or 10 µm sections were cut with either an RM2055 microtome (Leica) using a microtome knife, or a Shandon Finesse E+ microtome (Thermo) using a glass blade. Sections were stained with either double staining of 0.05% ruthenium red (Sigma-Aldrich) and toluidine blue (Sigma-Aldrich; 5s in each respectively, rinsed between staining's and afterwards with water), or 0.025% aqueus toluidine blue for 30s. Sections were mounted in either water or Histomount (National Diagnostics) and visualized either with a Leica 2500 Microscope or Zeiss Axioskop using 20x or 40x objectives.

For GUS-stained samples, Historesin was used. The GUS-staining protocol was adapted from (*47*). Samples were held in GUS-staining solution at 37°C until the appropriate staining level was reached prior to fixation.

<u>Fluorescent marker analysis, vibratome sections and EdU detection</u>

Lateral view of the fluorescent samples were analyzed in plates using Leica MZ165FC microscope, Hamamatsu C11440 digital camera, and Leica LAS X program.

For cross sections fluorescent samples were fixed as previously described (*3*), prior to embedding in 4% agarose. Agarose blocks were cut with the vibratome into 200 μm sections for confocal analysis. Sections were placed in PBS and stained with SR2200 (1:1000, Renaissance Chemicals) for cell wall staining. Lignified cell walls were stained with Basic Fuchsin, as previously described (*48*).

To visualize the EdU positive nuclei after EdU incubation, The Click-iT EdU Alexa Fluor 488 Imaging Kit (Thermo Fisher) was used for detection with a modified EdU detection mix (*49*). Samples were incubated in the detection mix for 1h and then transferred into PBS with SR2200 (1:1000).

<u>Confocal microscopy and image processing</u>

Confocal imaging was performed on PBS-mounted samples with a Stellaris 8 confocal microscope, except *35S:XVE>>gPLT5-TagRFP* analysis was carried out with Leica SP5 (20x and 63x objectives; Leica). Images were obtained using Las AF software (Leica). Samples visualized with multiple channels were imaged in the sequential scan mode. Confocal settings vary between experiments but were constant within experiments. The exception to this was cell wall staining to aid visualization. Here, SR2200 (cell wall) signal was adjusted during imaging (but not the fluorophore of interest) and therefore SR2200 settings varies between the sample and respective control.

<u>Image projections</u>

Circle unwrapping projections (**Fig. 3A**) were performed as previously described(*6*).

<u>Image analysis</u>

Image analysis and quantification were performed using Leica AF Lite 2.6.x, LithoGraphX 1.2.2 with Builder 1.2.2.7, and FIJI ImageJ v1.52(*50*). For **Fig.1D; and fig. S3A**, Cell numbers were calculated within 35 μm diameter area (primary xylem was in the center of this area). For image quantification in **Fig. 2E**, the distances of EdU-labelled nuclei were measured from the central point of each cross-section, as previously described(*3*). Locations of EdU-positive nuclei were expressed as a relative position along the radii of cross-sections. For each sample, the frequency distribution of EdU-labelled nuclei was then calculated and assigned into one of ten classes based on their relative positions in the cross-section. For each independent repeat, the mean and standard error of the frequency distributions of different cross-sections were calculated and plotted according to the treatments applied. To identify significant differences in the frequencies per distance class between the mock and induced conditions, Student's t-tests were applied.

For quantification of *plt3plt5plt7-tdna;IGE-ant* root cross-sections in comparison to wild type and *plt3plt5plt7-tdna* controls **(fig. S3D)**, sectors in the upper panel lacking secondary xylem vessel differentiation were considered as mutant sectors as such sectors were absent in controls. The lower panel considers secondary phloem differentiation around primary phloem pole. Controls developed secondary phloem around primary phloem pole, but sectors in *plt3plt5plt7-tdna;IGE-ant* plants with reduced secondary xylem formation produced also less secondary phloem (quantified as number of sieve elements) were considered to be mutant sectors.

For quantification of *gANT-3xYFP* fluorescence in **Fig. 3E**, only non-xylem-pole pericycle cell lineages were considered. The first, second or third nearest-neighboring cambial cell to vessels were defined as position 1, 2 or 3, respectively and assigned as *gANT-3xYFP*-positive or negative depending on the presence or absence of signal.

For quantification of *pANT:erRFP* fluorescence in **Fig. 4D**, the signal intensity in one cell file was measured from the outermost xylem vessel phloemward. For **Fig. 4H**, a line was drawn along the tangential axis of the root cross section between the outer edges of vessels. Cells between these two vessels on this line were considered to be at position zero. The radial cell file phloemward from this cell zero were marked as +1,+2,+3...+8; and towards xylem as -1,-2,-3.

<u>RT-qPCR</u>

11-days old plants with inducible *PXY* RNAi expression in *ANT* fluorescent reporter backgrounds (*35S:XVE>>PXY-RNAi; pANT:erRFP*) were transferred to 1/2GM plates containing either 5 µM of 17-β-estradiol (induced) or DMSO (mock) for three days. For RNA purification, 2 cm of primary roots 0.5 cm below the root-hypocotyl junction were harvested from ≥10 individuals for induced- or mock-treated plants. Total RNA from root samples was purified with RNeasy Plant Mini Kit (QIAGEN) with an on-column DNase I treatment (QIAGEN). cDNA was synthesized using the iScript™ cDNA Synthesis Kit (Bio-Rad) following the manufacturer's instructions. qRT-PCR experiments were carried out in 10 µl reaction volume using the LightCycler 480 SYBR Green I Master Mix (Roche Life Science) in a CFX384 Touch Real-Time PCR instrument (Bio-Rad). The PCR programme included an initial denaturation step at 95°C for 5 min, then 45 cycles of (95°C for 10s, 59°C for 10s, 72°C for 15s), followed by a melting curve analysis. Each sample was run three times. Expression levels were normalized using the comparative CT Method (ΔΔCT method) against the UBC21 reference gene expression (*51*). All primers used in qRT–PCR are listed in data S1.

<u>General methodology and statistical analysis</u>

The number of individual roots analysed is shown as *n* in the figures or figure legends, except for **Fig. 4, D and H**, *n* represents radial cell file. The fraction in the corners of some figures indicates the frequency of the observed phenotype. Before assessing statistical analyses, normality of residues distribution and variances homoscedasticity were checked using Shapiro's and Levene's tests, respectively, to determine the type of statistical analyses that can be used for each quantitative dataset. Accordingly, Wilcoxon-Mann-Whitney test was used to assess mean comparison for the **Fig. 2, C and G; Fig. 3, A and D; and Fig. 4, D and H** while student t-tests were used for the **Fig. 2E**. For multiple comparisons, Kruskal-Wallis and Dunn's post-hoc tests were used for the **Fig. 1, D and G**. For the **Fig. 3E**, categorical distribution was tested using Chi-square test. All the p-values for the different statistical comparison are available in the data supplemental information. Specific tests are detailed in the figure legends. All statistical analysis were performed in the R studio version 2023.06.0-421. For boxplot, the central line indicates the median; the bounds of the box show the 25th and 75th percentiles; and the whiskers indicate maximum and minimum values (the values out of the whiskers are outliers). For barplot, the bar height and the error bar represent the mean and the standard error of the mean, respectively, except **fig. S5F**, the error bar indicates standard deviation. For plots showing quantitative data, every individual data point was plotted on top of the plots. For the **Fig. 3A**, comparisons of the distance between P9A and TDIF conditions in both *pPLT5:erRFP* and *pPXY:erVenus* were obtained by sub-setting all the negative distances values (xylem region) present around the mean ± se of the fluorescence peak in P9A samples. *p-values* shown in the figure represent comparison of the mean distances between P9A and TDIF conditions using two tailed Wilcoxon-Mann-Whitney test.

**Modelling Methods**

Model network architecture and dynamics

In this paper we employ models of the radial patterning of the vasculature, using both a single cell and multicellular model setup. In the latter case, because of the rotational symmetry, we restrict ourselves to modeling vascular patterning in a single 1D cell file of the differentiating root vasculature. In our model all cells contain the same gene regulatory network describing the dynamics of auxin, TDIF, PXY, ANT, PLT (as a representative of PLT3, PLT5 and PLT7), and HD-ZIP III. Dynamics of the individual players making up this network are modeled using differential equations. Model cells can attain different vascular fates through experiencing different combinations of auxin and TDIF levels and through the regulatory network translating this into different gene expression patterns for PXY, ANT, PLT and HDZIPIII. Cells with sufficient PLT and ANT expression adopt cambial identity. Cells lacking these factors differentiate into xylem if they have sufficient HD-ZIP III, or phloem if they lack HD-ZIP III (for more details on the applied threshold levels to determine cell fate see later sections). In addition to intracellular expression dynamics, the TDIF signaling peptide is excreted and diffuses in the apoplast, while the ANT and PLT proteins diffuse between cells via plasmodesmata.

Gene regulatory and signaling network architecture

The architecture of the model gene regulatory and signaling network is shown in **(fig. S6)**. Yellow PE and orange TS symbols refer to supporting data the interaction was based on, and are summarized in the accompanying tables. In the below sections we describe the system of differential equations used to model the dynamics of the various network components.

Gene expression and signaling dynamics

PXY-TDIF expression and binding

At the heart of the model is the interaction between the auxin induced receptor protein PXY and the phloem produced mobile peptide ligand TDIF, that binds to it. We capture the production, movement, association, disassociation, and degradation of PXY and TDIF with the following set of equations.

$$\frac{dPXY_i}{dt} = p_{PXY}\frac{auxin_i^2}{auxin_i^2 + Km_{Aux,PXY}^2}\left((1 - fac_{PLTPXY}) + fac_{PLTPXY}\frac{Km_{PLT,PXY}^2}{PLT_i^2 + Km_{PLT,PXY}^2}\right) - d_{PXY}PXY_i - ass_{PXYTDIF}PXY_i * TDIF_i + diss_{PXYTDIF}PXYTDIF_i \quad (1)$$

$$\frac{dTDIF_i}{dt} = p_{TDIF} - ass_{PXYTDIF}PXY * TDIF_i + diss_{PXYTDIF}PXYTDIF_i - d_{TDIF}TDIF_i + TDIFdiffusion \quad (2)$$

$$\frac{dPXYTDIF_i}{dt} = ass_{PXYTDIF}PXY_i * TDIF_i - diss_{PXYTDIF}PXYTDIF_i - d_{PXYTDIF}PXYTDIF_i \quad (3)$$

, where i is an index for cell number, $prod_{PXY}$ is the maximum PXY production rate, $Km_{Aux,PXY}$ is the value for which the auxin-mediated induction of PXY is at half maximum, $fac_{PLTPXY}$ is the maximum fraction of PXY expression that can be repressed by PLT and $Km_{PLT,PXY}$ is PLT level at which this repression is at half maximum, $d_{PXY}$ is the PXY degradation rate, $ass_{PXYTDIF}$ and $diss_{PXYTDIF}$ are the rates of PXY-TDIF association and disassociation, $p_{TDIF}$ and $d_{TDIF}$ are maximum production and degradation rates for TDIF and $d_{PXYTDIF}$ is the degradation rate of PXY-TDIF. In the single cell layout, through varying the level of $p_{TDIF}$ we investigate the impact of TDIF level on cell fate. In the multicellular layout, $\mathrm{p_{TDIF}}$ equals 0 unless the cell is the predefined phloem cell that produces TDIF. Finally, the *TDIFdiffusion* term is 0 in the single cell layout. For further details on TDIF diffusion we refer to the subsection on protein diffusion in the section describing the multicellular model.

<u>HD-ZIP III expression</u>

Aside from inducing PXY, auxin also induces the expression of the xylem identity transcription factor HD-ZIP III. We use the following equation to describe HD-ZIP III expression dynamics:

$$\frac{dHDZIP_i}{dt} = p_{HDZIP}\frac{auxin_i^4}{auxin_i^4 + Km_{Aux,HDZIP}^4} - d_{HDZIP}HDZIP_i \quad (4)$$

, where $p_{HDZIP}$ is the maximum HD-ZIP III production rate, $Km_{AUX,HDZIP}$ is the auxin level at which HD-ZIP III production is half maximum and $d_{HDZIP}$ is the degradation rate of HD-ZIP III. Note the use of a Hill-coefficient of 4 as compared to the Hill-coefficient of 2 in our other equations. This higher Hill-coefficient results in a slightly more switch-like dependence of auxin levels which ensures that the high-auxin requiring (high Km) HD-ZIP III can become more highly expressed and hence dominant over the lower-auxin requiring (low Km) PXY at high auxin levels, ensuring a stable xylem domain.

<u>ANT and PLT expression</u>

ANT expression is induced by both auxin and PXY-TDIF, while being repressed by HD-ZIP III. We assume that auxin and PXY-TDIF induction of ANT function additively, and are independently antagonized by HD-ZIP III. Combined this results in the following equations:

$$\frac{dANT_i}{dt} = p_{ANT}\left(\frac{act_{Aux,ANT} * auxin_i^2}{auxin_i^2 + Km_{Aux,ANT}^2} + \frac{act_{Aux,PXY} * PXYTDIF_i^2}{PXYTDIF_i^2 + Km_{PXY,ANT}^2}\right) - d_{ANT}ANT_i + ANTdiffusion$$

$$act_{Aux,ANT} = max_{Aux,ANT}\left(\left(1 - rep_{HDZIP,Aux}\right) + \frac{rep_{HDZIP,Aux} * Km^2_{HDZIP,ANT}}{HDZIP_i^2 + Km^2_{HDZIP,ANT}}\right) \tag{5}$$

$$act_{PXY,ANT} = max_{PXY,ANT}\left(\left(1 - rep_{HDZIP,PXY}\right) + \frac{rep_{HDZIP,PXY} * Km^2_{HDZIP,ANT}}{HDZIP_i^2 + Km^2_{HDZIP,ANT}}\right)$$

, where $p_{ANT}$ is the maximum ANT production rate, $act_{Aux,ANT}$ and $act_{PXY,ANT}$ are the auxin and PXY-TDIF dependent ANT induction functions that depend on HDZIP-III repression, $Km_{Aux,Ant}$ and $Km_{PXYANT}$ are the values of auxin and PXY for which their activation of ANT expression are at half maximum, $d_{ANT}$ is the degradation rate of ANT, $max_{Aux,ANT.}$ and $max_{PXY,,ANT}$ are the maximum fractions of auxin and TDIF signalling mediated ANT induction, $rep_{HDZIP,Aux}$ and $rep_{HDZIP,PXY}$ are the maximum fractions of these that can be repressed by HDZIP-III, and $Km_{HDZIP,ANT}$ is the HDZIP-III level at which this repression is half maximal. *ANTdiffusion* is 0 in the single cell layout, and further defined in the subsection on protein diffusion.

In contrast to ANT, PLT expression is only induced by TDIF peptide signaling and is not repressed by HD-ZIP III, resulting in the following simpler equation:

$$\frac{dPLT_i}{dt} = p_{PLT}\frac{PXYTDIF_i^2}{PXYTDIF_i^2 + Km^2_{PXY,PLT}} - d_{PLT}PLT_i + PLTdiffusion \tag{6}$$

, where $p_{PLT}$ is the maximum PLT expression rate, $Km_{PXY,PLT}$ is the PXY-TDIF level for which the expression of PLT is half maximum, and $d_{PLT}$ is the degradation rate of PLT. *PLTdiffusion* is 0 in the single cell layout, and further defined in the subsection on protein diffusion section.

<u>Model parametrization</u>

After defining network architecture based on experimental data, and establishing the basic differential equations-based model describing the dynamical interactions resulting from this network architecture, we next need to determine the (range of) parameter values to be used for the model.

<u>Defining a parameter range</u>

<u>Production and degradation rates</u>

As a first step, in absence of quantitative data, we scale the maximum level for the signaling molecules auxin and TDIF, as well as for the transcription factors HD-ZIP III, ANT, and PLT and the receptor protein PXY to a value of 100, which can be interpreted as meaning 100%. The maximum level of 100 implies that the ratio of production over degradation rates for each gene is set at 100, while the actual rates can be varied. We typically use a degradation rate of $0.0002s^{-1}$ and hence a production rate of $0.02[]s^{-1}$ (**see table S1**), which albeit somewhat arbitrary is chosen such that it is significantly faster than the rate of cell division, ensuring that despite growth and division driven dilution of proteins, concentrations are always near steady state. While in the current model there is no growth and division of cells, this allows seamless future incorporation of these processes without requiring reparametrization.

As an exception to the above, for the receptor protein PXY we applied a higher degradation rate (typically a degradation rate of $0.0012s^{-1}$ is used, see **table S1**). The rationale behind this higher degradation is that this encompasses both receptor internalization and turnover. Assuming that upon binding of the TDIF ligand to the PXY receptor, the PXY-TDIF complex turns over at the same rate as isolated PXY, this enables us to investigate the effect of receptor mediated TDIF sequestration and enhanced degradation on TDIF signaling gradient formation. In the parameter sweep the $p_{TDIF}$ parameter is varied between 0 and 0.028.

<u>Km values and ANT expression regulation</u>

As a second step, having set the maximum level of 100, we are now able to deduce the relevant range for model Km values. Put simply, Km values of above 100 would cause downstream activated regulatory factors to never reach their maximum levels or downstream repressed factors to never approach their minimum levels, whereas Km values of 5 or lower would nearly abolish the dependence on an upstream regulatory factor, setting a first broad range of relevant values. Based on the biological data, and the simulated auxin and TDIF profiles in our model, reasonable Km values can subsequently be further constrained.

First, given that the data show that at the phloem side, where the lowest auxin levels occur (parametrized to a value of 8 in our model, either imposed in single cell settings or resulting from auxin gradient dynamics, see multicellular model section) none of the auxin-induced factors (HD-ZIP III, ANT, PXY) are expressed, the Km of these factors for auxin induction must be at least 15. Additionally, since these factors are known to be highly expressed at the xylem side of the vasculature where auxin levels are highest (up to 100 in our model), the maximum value for these Km values should not be higher than 50 (70 for $Km_{Auxin,HDZIP}$ because of its higher Hill coefficient). Taking into account the observation that HD-ZIP III is expressed exclusively in the xylem, where auxin levels are highest, while PXY is expressed in both xylem and cambium and ANT is only expressed in the cambium, where auxin levels are intermediate, we can further constrain and order the Km values. (Note that ANT expression in the xylem is antagonized by HD-ZIP III, see **Fig. 3E**). Specifically, these data imply that $Km_{Auxin,HDZIPIII}$ is larger than $Km_{Auxin,PXY}$ and $Km_{Auxin,ANT}$. Finally we constrain $Km_{HDZIP,ANT}$ to only become half activated at an HD-ZIP III level of 30 to restrict HD-ZIP III activity to the high auxin domain. Thus, we have established a biologically plausible range for the 4 Kms in **table S2**.

The last four parameters in **table S2** relate to the induction of ANT by auxin and PXY and its repression by HD-ZIP III. For these we again use biological data and practical considerations to constrain their values. Since auxin also induces PXY, the additive induction of ANT by auxin and PXY corresponds to a direct activation and indirect activation by auxin. To constrain maximum ANT expression in our parameter sweep these two fractions were varied in an anti-correlated manner, i.e. if one was increased the other was decreased proportionally. Since the PXY-TDIF interaction is known to be critical for ANT expression, we set the maximum direct auxin fraction for *ANT* induction to 0.4 and hence the minimum for the indirect PXY-TDIF induction of ANT to 0.6. The last two parameters, $rep_{HDZIP,Aux}$ and $rep_{HDZIP,PXY}$, represent the extent to which HD-ZIP III represses the direct auxin induction of ANT, compared to the indirect PXY-TDIF induction of ANT and are independently varied in our parameter sweep. A minimum of 0.4 was set as HD-ZIP III as the biological data dicttes that HDZIP-III at least partially represses ANT induction and we are not interested in the regime where this does not occur for the parameter sweep.

<u>PXY-TDIF signaling related parameters</u>

The parameters in **table S3** refer to the translation of PXY-TDIF signal to PLT expression as well as the PLT repression of PXY expression. Since PXY-TDIF is formed at the region of overlap between opposing PXY and TDIF gradients, where both PXY and TDIF levels are substantially submaximal we reasoned that as a minimum requirement parameter settings should result in maximum TDIF and PXY levels (100) translating into a high level of PXY-TDIF complex (e.g. 80) and little PXY and TDIF to remain unbound (20). This gives us the following constraint for PXY-TDIF association and dissociation rates:

$$\frac{diss_{PXYTDIF}}{ass_{PXYTDIF}} = \frac{[PXY] * [TDIF]}{[PXYTDIF]} = \frac{20 * 20}{80} = 5 \qquad (7)$$

Additionally, we assume that association and dissociation rates are faster than the turnover of these proteins themselves. We settled on 0.02 for the association rate (and thus 0.1 for the dissociation rate), corresponding to 100 times the degradation rate of PXY.

In practice PXY-TDIF binding only occurs at intermediate PXY and TDIF values. Thus, intermediate values of PXY-TDIF complex should be capable of inducing the high levels of ANT and PLT expression required for cambial identity, while low PXY-TDIF levels should not lead to their expression. Combined this sets a range of Km values for PXY-TDIF induced expression for ANT between 15 and 35, and for PLT between 20 and 40. We assigned slightly lower values for $KM_{PXYTDIF,ANT}$ than for $KM_{PXYTDIF,PLT}$ for two reasons. First, PLT represses PXY, so to somewhat protect PXY from this, PLT induction should require significant levels of PXY-TDIF. Secondly, HD-ZIP III represses ANT, so to compensate for this, a slightly lower PXY-TDIF should already sufficiently induce ANT.

The repression of PXY by PLT results in a negative feedback loop that puts a cap on the PXY-TDIF induced *PLT* and *ANT* expression. To still enable the high *PLT* and *ANT* expression observed experimentally, we assume that PLT can maximally reduce PXY levels by 30%, and that for this maximum repressive activity high PLT levels (Km>30) are needed. We speculate that such a parametrization may serve *in planta* as a sort of homeostatic mechanism, enabling cells to generate high expression of *PLT* and *ANT* with moderate PXY-TDIF levels, while preventing even higher PLT and ANT levels when PXY-TDIF levels further increase. Since the maximum and Km of PLT mediated PXY repression have similar effects, we keep this maximum repression constant while varying the Km to vary overall repression in the performed parameter sweep.

Cell fate threshold values

Finally, we need to set the values for the threshold parameters determining how we translate gene expression patterns into vascular cell fate. Based on our experimental data it is the activity of PLT and ANT that induce cambium identity. Given that PLT and ANT individually have a maximum protein level of 100, a threshold level above 100 implies that cambium identity requires both factors being present in significant amounts. Since knockout studies suggest this not to be the case (*20*), we chose a value of 75 for the cambium identity threshold.

Similarly, experimental data indicates that HD-ZIP III expression induces xylem cell fate. We set the threshold value for HD-ZIP III above which xylem fate is induced to 30. Note that the precise level of this threshold mainly determines the auxin level required to shift from phloem to xylem fate. Phloem fate occurs if neither the demands for xylem nor cambium fate have been met by the cells expression state, thus PLT+ANT <75 and HDZ-IP III<30.

Parameter sweep

To determine the robustness with which the above network reproduces the biological observation of high auxin levels resulting in xylem differentiation, high TDIF and low auxin levels resulting in phloem fate, and intermediate values inducing cambial identity, we apply a parameter sweep across the previously identified plausible ranges of parameter values. We vary each of the parameters from **tables S2 and table S3** in set increments between these extreme values. We perform this parameter sweep for the single cell model, exposing it for each investigated parameter setting to a range of auxin and TDIF levels between 0 and 100. For each pair of auxin and TDIF levels investigated, we score across the entire range of the parameter sweep the frequency with which the cell converges to the different possible cell types, enabling us to draw 2D auxin-TDIF cell fate maps. As long as the combined threshold for PLT+ANT for cambium formation remains below 100, there is only a quantitative shift in model outcome (**fig. S7, A** and **B**), while the general behavior remains robust.

Role of HD-ZIP III in specifying xylem

By taking specific subsets of the parameter sweep, where we keep one parameter constant, we can directly compare the overall effect of that parameter by contrasting two extreme values. While for most parameters we set relatively narrow ranges, the HD-ZIP III repression of ANT via $rep_{HDZIP,Aux}$ and $rep_{HDZIP,PXY}$ were left free to vary between barely repressing ANT at 0.4, to completely repressing ANT at 1.0. By zooming in on a subset of the parameter set we show how a strong repression of ANT by HD-ZIP III is able to shift cells with high auxin and intermediate TDIF from cambial identity to xylem identity (compare **fig. S7, A** and **C**). Thus, this strong HD-ZIP III activity can safeguard the high auxin xylem from intermediate TDIF mediated conversion to cambium.

Extension to multicellular model

In the multicellular model we created a 1D tissue strand with a xylem organizer cell on the left, a mature phloem cell on the right, and 1-3 cambial cells in between. On this 1D cell file we superimpose an auxin gradient which has its maximum at the xylem organizer cell, and incorporate production of TDIF occurring in the mature phloem cells (see Eq. 2). The model was run till it reached steady state before analyzing outcomes. To take into account cell size differences, we applied for the different cell types the following cell widths: Xylem cell 12 µm, cambium 4 um and phloem 8 µm. The height was set at 25 µm, which is somewhat arbitrary due to the 1D nature of the model. Model simulations use the rate parameters of **table S1**, and the default auxin and PXY-TDIF parameter values of **table S2 and tableS3** unless explicitly stated differently, as described in **table S5 and table S6.**

<u>Superimposed auxin gradient</u>

Auxin is a key player in cambium development, being a major regulator for HD-ZIP III, PXY and ANT expression. In the cambium, experimental data show a characteristic auxin gradient with its maximum at the most cambium-ward adult xylem cell that gradually decreases towards the phloem. In absence of sufficient data on the relative importance of longitudinal and transversal auxin transport and local auxin production in shaping this gradient, instead of explicitly modeling auxin dynamics, we superimpose an auxin gradient according to the following equation:

$$auxin_{xylem} = max_{auxin}$$
$$auxin_1 = \left(max_{auxin} - drop_{xylem}\right) \qquad (8)$$
$$auxin_{i>1} = \frac{\left(max_{auxin} - drop_{xylem}\right)}{mod_{cambium} * i^2}$$

,where i is the cell number starting at 1 in the most xylem-ward cambial cell, $max_{auxin}$ is the level of auxin at the auxin maximum in the xylem (default level 100), $drop_{xylem}$ is the initial drop relative to the maximum for the first cambial cell (default level 60), and $mod_{cambium}$ cambial modulates the further reduction of auxin as distance (measured in number of cells) from the xylem increases (default level 1.25). Overall this results in a semi-exponentially decreasing auxin gradient.

<u>Dynamic protein diffusion</u>

Apoplastic diffusion of the phloem secreted TDIF peptide into the cambium and towards the xylem is critical for achieving TDIF signalling. In the model we parametrized TDIF production and diffusion such that under standard parameter conditions, TDIF could reach 3 cells beyond the phloem at a sufficient level for it to induce cambial identity.

The transcription factors ANT and PLT are capable of moving between cells through plasmodesmata (*19*), and thus display cytoplasmic diffusion. Based on the difference in size between small TDIF peptides and full blown ANT and PLT proteins and the different nature of their diffusion (apoplastic versus symplastic) we chose ANT and PLT diffusion rates to be an order of magnitude lower than that of TDIF (see **table S4**).

In all 3 cases, to ensure mass balance when updating concentrations for diffusional exchange between different sized cells, we apply a size dependent scaling.

$$TDIFdiffusion_i = \frac{D_{TDIF}}{size_i} * (TDIF_{i-1} + TDIF_{i+1} - 2 * TDIF_i)$$
$$ANTdiffusion_i = \frac{D_{ANT}}{size_i} * (ANT_{i-1} + ANT_{i+1} - 2 * ANT_i) \qquad (9)$$
$$PLTdiffusion_i = \frac{D_{PLT}}{size_i} * (PLT_{i-1} + PLT_{i+1} - 2 * PLT_i)$$

HD-ZIP III repression vs sequestering

In **(fig. S8; and Fig. 4)** of the main manuscript we compare two alternative parameter regimes. Specifically, we used either the strong HD-ZIP III' repression of ANT **(Fig. 4, A and E; fig. S8B)** or we used the "strong sequestering" parameter values (**Fig. 4, B and F; fig. S8C**). Under these latter settings HD-ZIP III does not repress the PXY-TDIF based expression of ANT at all, and instead exclusively represses the auxin fraction. To further enhance the possibility for strong sequestration, we lowered the $diss_{PXYTDIF}$ from 0.1 to 0.02.

To investigate the relevance of sequestration for model outcomes we performed the strong HD-ZIP III simulations in the spatial 1D model while omitting sequestration. We achieve this in an artificial manner, by using the association and dissociation rates to compute PXY-TDIF complex levels as an input for ANT and PLT expression (Eq. 10 below) , yet not applying the association and dissociation dynamics in Eqs 2 and 4, and not using Eq. 3., thereby not diminishing free TDIF (and PXY) levels through binding to PXY. As a consequence, TDIF tissue motility is not limited due to PXY binding.

$$ass * (PXY_{tot} - X) * (TDIF_{tot} - X) = diss * X$$
$$PXY_{tot} * TDIF_{tot} - (PXY_{tot} + TDIF_{tot})X + X^2 = \frac{diss}{ass} * X \qquad (10)$$
$$PXY_{tot} * TIDF_{tot} - \left(PXY_{tot} + TDIF_{tot} + \frac{diss}{ass}\right) * X + X^2 = 0$$
$$X = \frac{\left(PXY_{tot} + TDIF_{tot} + \frac{diss}{ass}\right) - \sqrt{\left(PXY_{tot} + TDIF_{tot} + \frac{diss}{ass}\right)^2 - 4 * (PXY_{tot}TDIF_{tot})}}{2}$$

with X the computed PXY-TDIF complex level.

Note that a similar effect of strongly reduced TDIF sequestration can be achieved through reducing the binding of TDIF to PXY, while simultaneously increasing the effectiveness of PXY-TDIF mediated ANT and PLT expression.

This way, in one subset sequestration effects on motility were removed while HDZIPIII strongly represses ANT expression, while in the other subset of simulations sequestration effects were enhanced while the HDZIPIII repression was lowered.

Variable gradients

To investigate the capacity of the network to robustly integrate and respond to various auxin and TDIF signaling inputs, we varied the auxin and TDIF production levels in a 3 cells wide cambium (5 cells in total) **(fig. S9)**. In this larger cambium there is more room for differences in spatial overlap of gradients for different auxin gradient and TDIF production and diffusion parameter settings (**table S6**). These simulations used the strong sequestration parameters from **table S5**. We ascribe cell fates to the cells based on the cell fate threshold values section as described above.

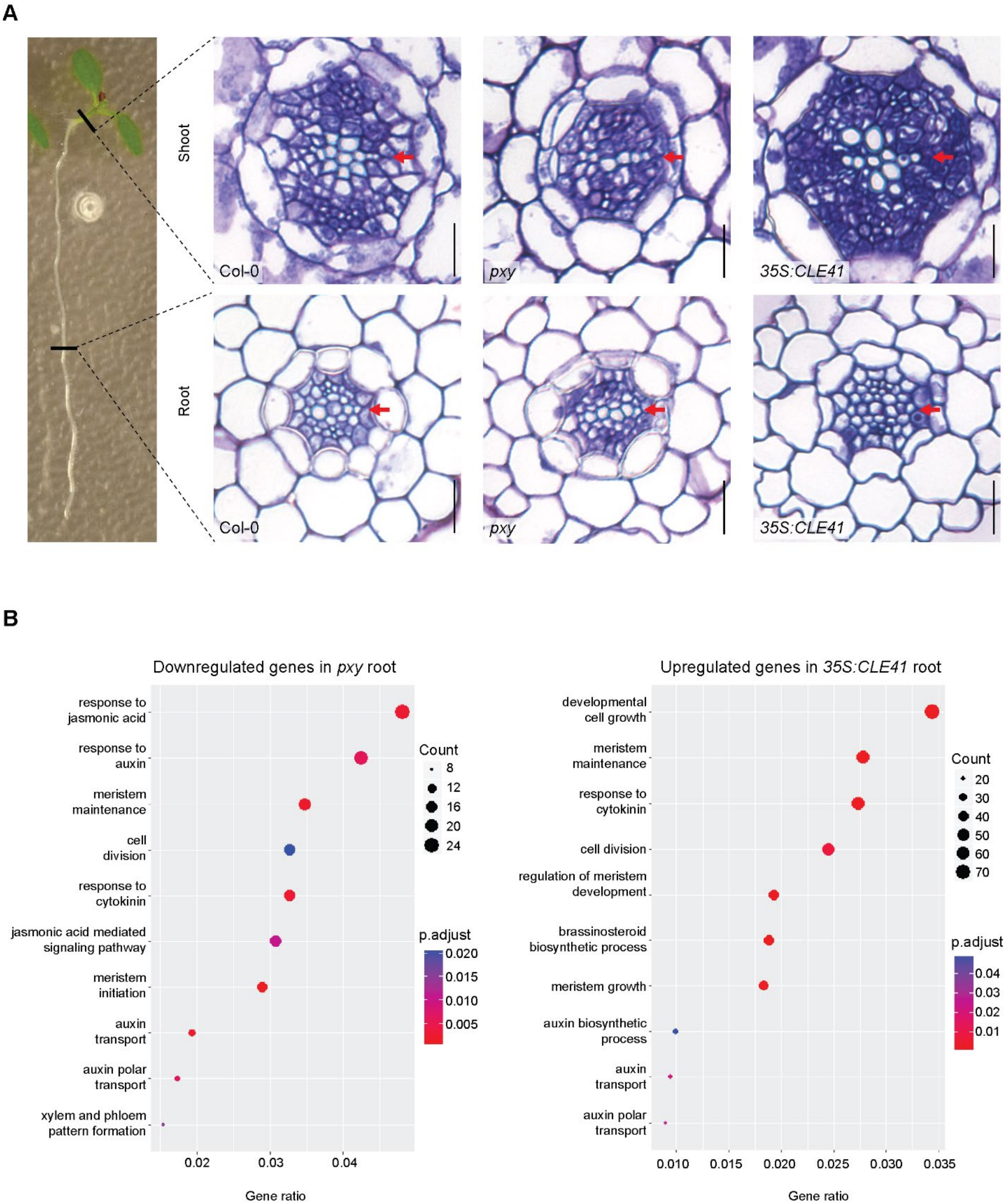
A
Shoot
Root
Col-0
pxy
35S:CLE41
Col-0
pxy
35S:CLE41
B
Downregulated genes in pxy root
response to jasmonic acid
response to auxin
meristem maintenance
cell division
response to cytokinin
jasmonic acid mediated signaling pathway
meristem initiation
auxin transport
auxin polar transport
xylem and phloem pattern formation
Count
8
12
16
20
24
p.adjust
0.020
0.015
0.010
0.005
0.02
0.03
0.04
Gene ratio
Upregulated genes in 35S:CLE41 root
developmental cell growth
meristem maintenance
response to cytokinin
cell division
regulation of meristem development
brassinosteroid biosynthetic process
meristem growth
auxin biosynthetic process
auxin transport
auxin polar transport
Count
20
30
40
50
60
70
p.adjust
0.04
0.03
0.02
0.01
0.010
0.015
0.020
0.025
0.030
0.035
Gene ratio

**Fig. S1. Transcriptomic analysis of *pxy* and CLE41 overexpression**

**(A)** Left, an image of a seedling showing the position of cross section shown on the right. Cross-sections of 7-day old Wild type Col-0, *pxy, 35S:CLE41* of hypocotyls and roots. **(B)** Gene Ontology term of reduced *pxy* and enriched in *35S:CLE41* root respectively. Red arrows mark the primary xylem axis. Scale bars 20 µm.

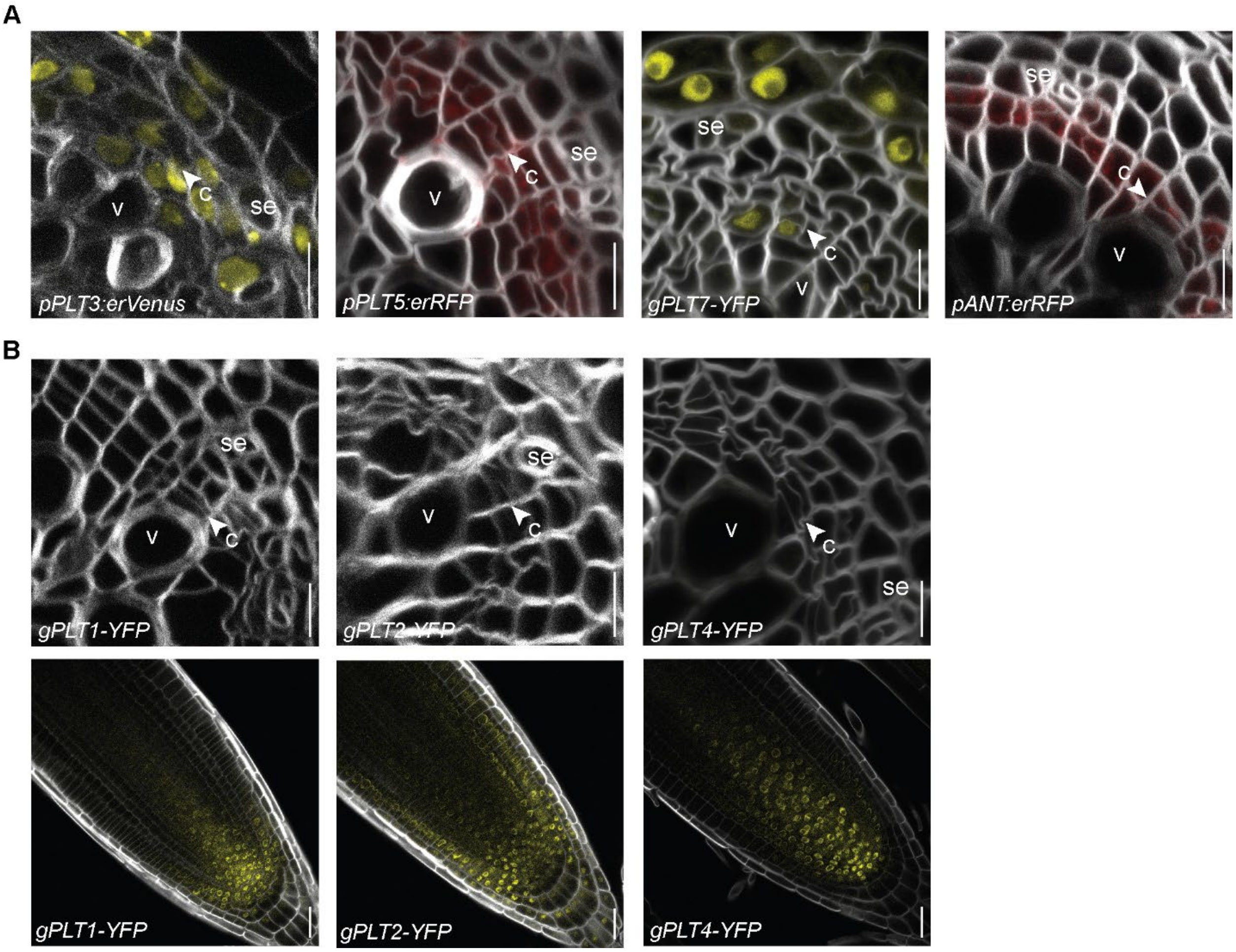
A
B
se
c
v
pPLT3:erVenus
pPLT5:erRFP
gPLT7-YFP
pANT:erRFP
gPLT1-YFP
gPLT2-YFP
gPLT4-YFP

**Fig. S2. Four *AIL*/*PLT*s are expressed in root cambium**

**(A)** Confocal cross-sections of 14-day-old *pPLT3:erVenus, pPLT5:erRFP, gPLT7-YFP* and *pANT:erRFP* roots. **(B)** Confocal cross-sections of 14-day-old roots and longitudinal view of root tips of *gPLT1-YFP, gPLT2-YFP* and *gPLT4-YFP* show expression in root tip as previously reported(*19*), however they show no fluorescence in root vascular cambium. White arrowheads mark recent cell division. Vessels (v), cambium (c), sieve element (se). Scale bars 10 µm.

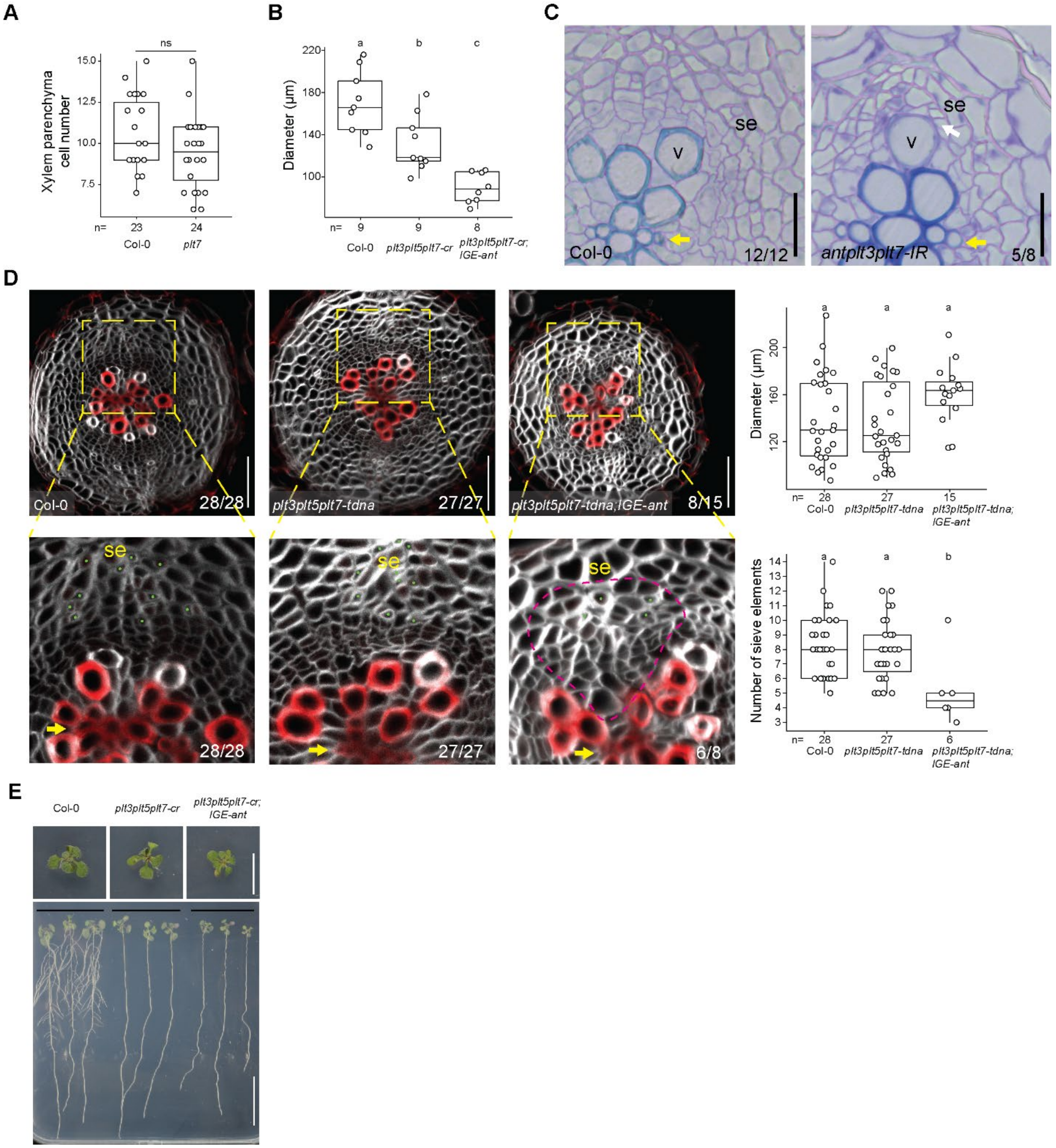

A
ns
Xylem parenchyma cell number
15.0
12.5
10.0
7.5
n= 23 24
Col-0
plt7
B
a b c
Diameter (µm)
220
180
140
100
n= 9 9 8
Col-0
plt3plt5plt7-cr
plt3plt5plt7-cr; IGE-ant
C
se
v
Col-0
12/12
se
v
antplt3plt7-IR
5/8
D
Col-0
28/28
plt3plt5plt7-tdna
27/27
plt3plt5plt7-tdna;IGE-ant
8/15
se
28/28
se
27/27
se
6/8
a a a
Diameter (µm)
200
160
120
n= 28 27 15
Col-0
plt3plt5plt7-tdna
plt3plt5plt7-tdna; IGE-ant
a a b
Number of sieve elements
14
13
12
11
10
9
8
7
6
5
4
3
n= 28 27 6
Col-0
plt3plt5plt7-tdna
plt3plt5plt7-tdna; IGE-ant
E
Col-0
plt3plt5plt7-cr
plt3plt5plt7-cr; IGE-ant

**Fig. S3. Histological analysis of *ail/plt* mutants**

**(A)** Boxplot showing xylem parenchyma cell number in Col-0 and *plt7*. **(B)** Quantification of vascular diameter from Fig. 1F. **(C)** Cross section of 14-day-old Col-0 and an *antplt3plt7* triple mutant (*ant-4ail6-2ail7-IR*)(*16*). **(D)** Confocal cross-section of 13-days-old roots of Col-0, *plt3plt5plt7-tdna, plt3plt5plt7-tdna;IGE-ant* and the quantification of the vascular diameter and number of sieve elements (right panels). 8/15 of *plt3plt5plt7-tdna;IGE-ant* roots showed sectors without vessel production. 6/8 of these sectors occurred in the position of phloem pole, and these 6 sectors were used in quantification of sieve elements (bottom right panel). Inset images show reduced number of sieve elements (marked with green dot) and the sector marked with dotted line (magenta). Cell wall stained with SR2200 (grey), lignified cell walls are stained with 0.1% basic fuchsin (red). **(E)** Gross morphology of 11-day-old plants. Significance difference was tested by Student t-test in (A) (ns = $p>0.05$). Letters indicate significant differences using one-way ANOVA with a Tukey post hoc test in (B), or using Kruskal-Wallis with Dunn post hoc test in (D). Numbers in (C) and (D) represent the frequency of the observed phenotypes. Yellow arrows mark the primary xylem axis. White arrow marks the position of sieve element touching vessel. Vessels (v), sieve elements (se), Scalebars 20 µm (C), 50 µm (D) and 1 cm (E).

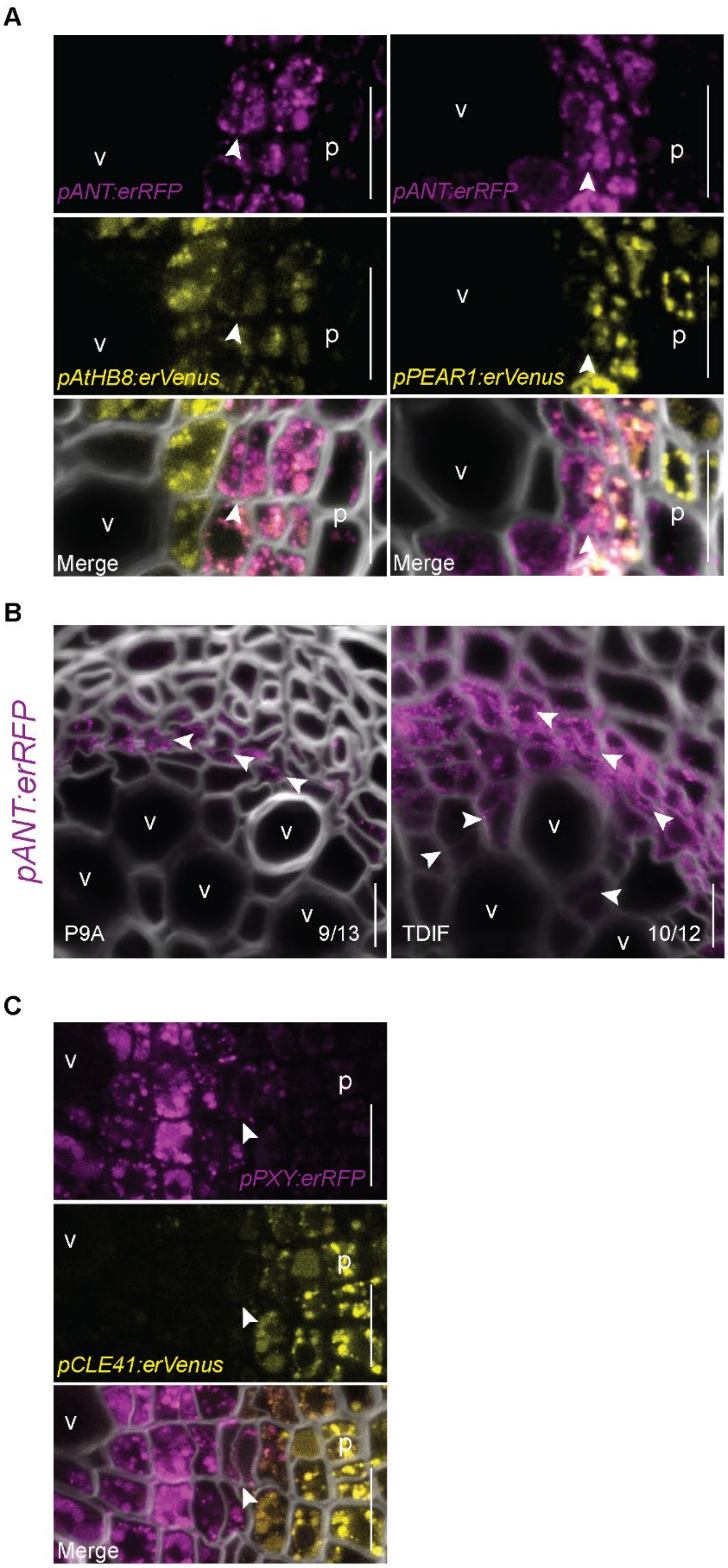
A
pANT:erRFP
pANT:erRFP
pAtHB8:erVenus
pPEAR1:erVenus
Merge
Merge
v
p
B
pANT:erRFP
P9A
9/13
TDIF
10/12
C
pPXY:erRFP
pCLE41:erVenus
Merge

**Fig. S4. Expression patterns of key cambial regulators**

**(A)** Confocal cross-sections of 12-day-old roots expressing *ANT* double markers. **(B)** Confocal cross-sections of *pANT:erRFP* after 1-day TDIF treatment in 14-day-old plants. **(C)** Confocal cross-sections of 14-day-old *pPXY:erRFP;pCLE41:erVenus*. Numbers in (B) represent the frequency of the observed phenotypes. White arrowheads mark recent cell division. Phloem (p). Vessels (v). Scale bars 10 µm (A to C).

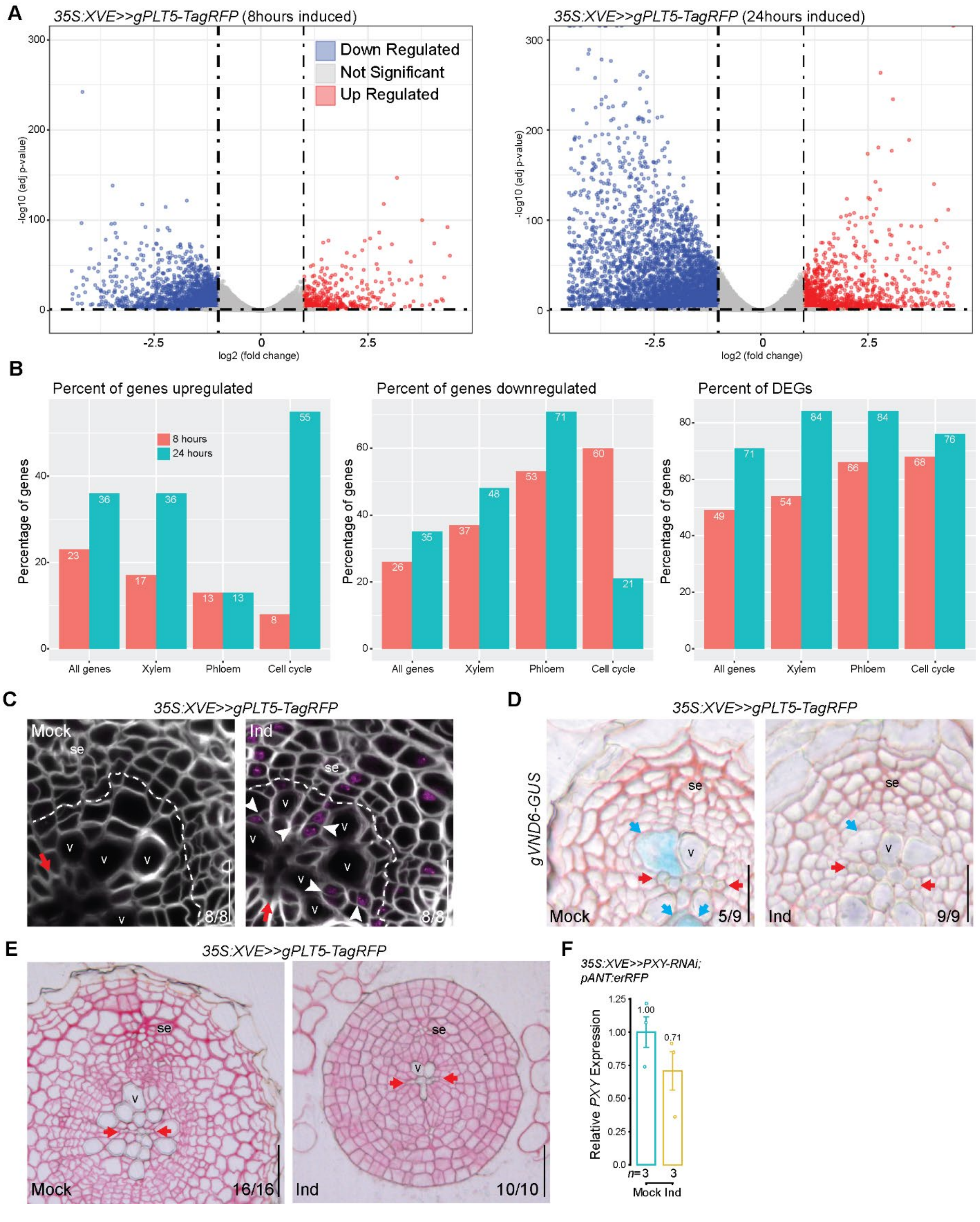
A
35S:XVE>>gPLT5-TagRFP (8hours induced)
Down Regulated
Not Significant
Up Regulated
-log10 (adj p-value)
log2 (fold change)
35S:XVE>>gPLT5-TagRFP (24hours induced)
B
Percent of genes upregulated
Percent of genes downregulated
Percent of DEGs
8 hours
24 hours
Percentage of genes
All genes
Xylem
Phloem
Cell cycle
C
35S:XVE>>gPLT5-TagRFP
Mock
Ind
se
v
8/8
D
35S:XVE>>gPLT5-TagRFP
gVND6-GUS
5/9
9/9
E
35S:XVE>>gPLT5-TagRFP
16/16
10/10
F
35S:XVE>>PXY-RNAi; pANT:erRFP
Relative PXY Expression
1.00
0.71
n=3
3
Mock Ind

**Fig. S5. Analysis of the consequences of *PLT5* overexpression**

**(A)** Volcano plot of all the transcript after 8 hours or 24 hours induction of *35S:XVE>>PLT5-TagRFP* RNA-seq. The dotted horizontal line corresponds to a Benjamini–Hochberg corrected significance of Padj Value <0.05. The dotted vertical lines bound the minimal fold-change for the most-differentially-expressed genes. **(B),** Bar plot shows the percentage of the genes that are upregulated, downregulated and differentially expressed genes (DEGs) in categories of All genes, xylem, phloem and cell cycle genes for 8 and 24 hours using PValue <0.05. **(C),** Confocal cross-section of *35S:XVE>>PLT5-TagRFP* after 2-day induction (in 12-day-old plants) showing ectopic cell division in xylem parenchyma. (Mock: No cell division in xylem; Ind: Cell divisions in xylem) most recent cell divisions in cambium marked with dotted lines (white). **(D),** Cross-section of *35S:XVE>>PLT5-TagRFP;gVND6-GUS* after 2-day induction (in 8-day-old plants). **(E),** Cross-section of *35S:XVE>>PLT5-TagRFP* after 7-day induction (in 8-day-old plants). **(F),** RT-qPCR showing the reduced *PXY* expression in *35S:XVE>>PXY-RNAi;pANT:erRFP*. Barplot shows average with ± sd. Numbers in panels (C to E) represents the frequency of the observed phenotypes. White arrowheads mark recent cell division, red arrows mark the primary xylem axis, and blue arrows marks the expanding xylem vessels. Vessels (v). Sieve elements (se). Scale bars 20 µm (C to E).

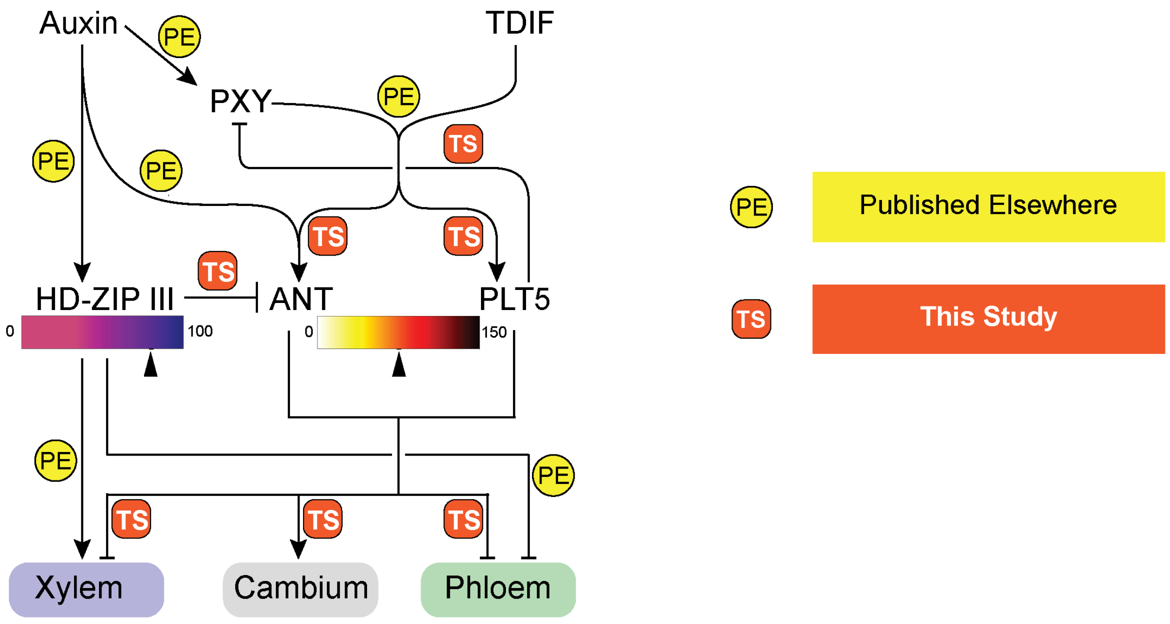


| | | |
|---|---|---|
| Published Elsewhere | Auxin ->PXY | Smetana et al., 2019 |
| | PXY & TDIF | Hirakawa et al., 2008; Etchells et al., 2010; Morita et al., 2016 |
| | Auxin -> HD-ZIP III | Smetana et al., 2019; Donner et al., 2009; Ursache et al., 2014 |
| | Auxin -> ANT | Smetana et al., 2019; Yamaguchi et al., 2013 |
| | HD-ZIP III -> Xylem | Zhong and Ye, 1999; Smetana et al., 2019; Ohashi-Ito et al., 2005; Carlsbecker et al., 2010 |
| | HD-ZIP III -\| Phloem | Smetana et al., 2019; Miyashima et al., 2019 |

| | | |
|---|---|---|
| **This Study** | PXY-TDIF -> ANT | Fig. 1B; fig. S4B |
| | PXY-TDIF -> PLT5 | Fig. 1B; Fig. 3A |
| | PLT5 -\| PXY | Fig. 2B; Fig. 3D |
| | HD-ZIP III -\| ANT | Fig. 3E |
| | ANT/PLT5 -\| Xylem | Fig. 2, F and G; fig. S5E |
| | ANT/PLT5 -> Cambium | Fig. 2, D and E; fig. S5C |
| | ANT/PLT5 -\| Phloem | Fig. 2, F and G; fig. S5E |

**Fig. S6. Signaling network in vascular cambium used in the modelling**

Top panel indicates the same network of modelled regulatory interactions driving cambial cell fate decision making as shown in Fig. 3C. Bottom panels indicate the experimental support for the different incorporated regulatory interactions categorized according to being published elsewhere (yellow, PE) or observed in this current study (orange, TS).

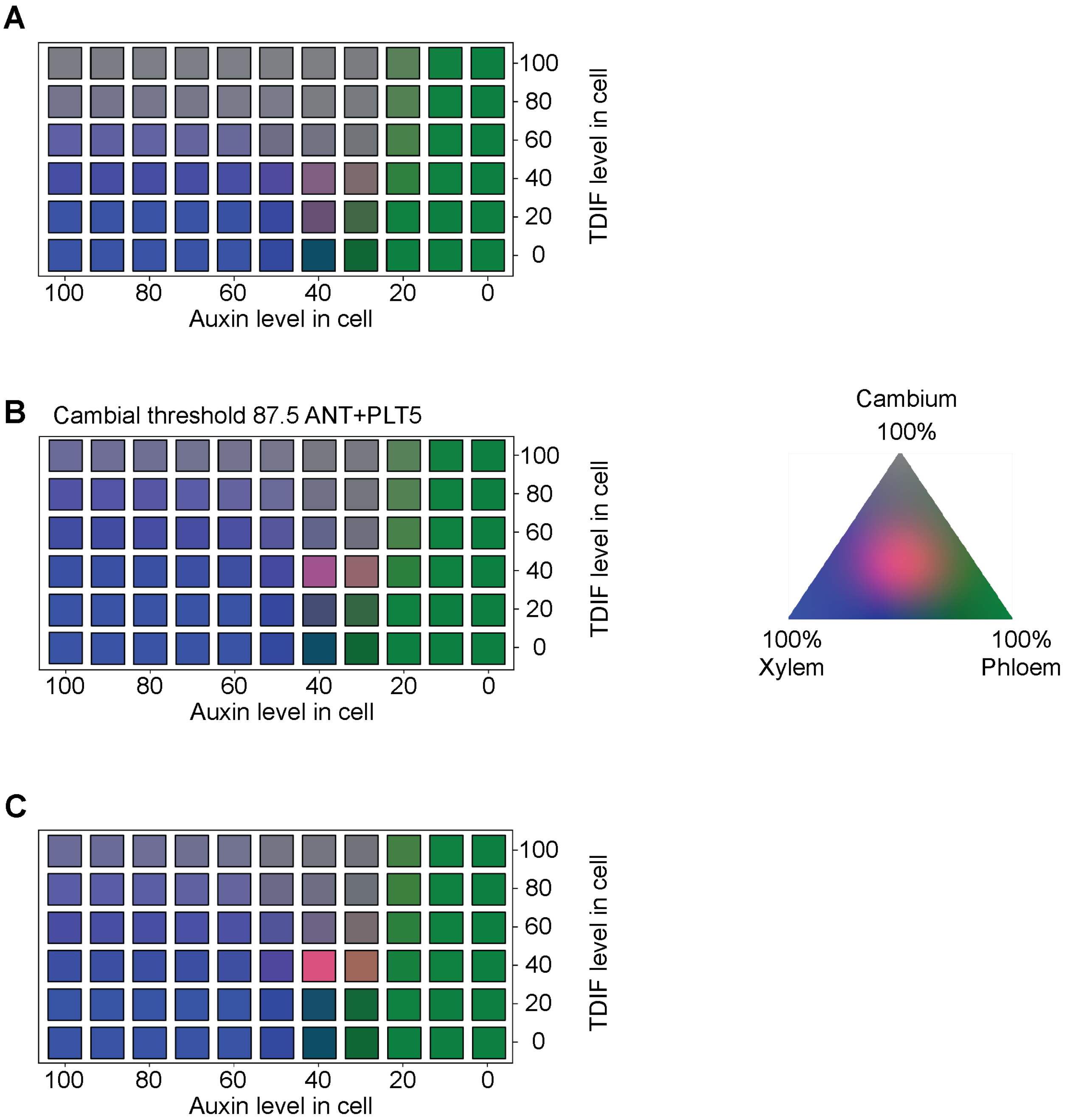
A
B
Cambial threshold 87.5 ANT+PLT5
C
100
80
60
40
20
0
TDIF level in cell
Auxin level in cell
Cambium
100%
100%
Xylem
100%
Phloem

**Fig. S7. Fate map for large scale parameter sweeps**

**(A)** Fate map of overall parameter sweep including both strong and weak HD-ZIP III mediated ANT repression for a cambial fate threshold of 75. For the range and sampling interval of parameter values used for the parameter sweep see Supplementary Computational Methods. Each Auxin-TDIF combination is colored according to the fraction of simulations that acquire xylem, cambium, or phloem identity according to the color triangle. **(B)** Alternative fate map of an overall parameter sweep using a cambial fate threshold of 87.5. Raising this threshold to 87.5 reduces the cambial domain but retains the same qualitative behavior as shown in (A). **(C)** Fate map for a parameter sweep constrained to strong HD-ZIP III mediated ANT repression (see Modelling Methods), showing an expansion of xylem over cambial fate for high auxin and high TDIF levels.

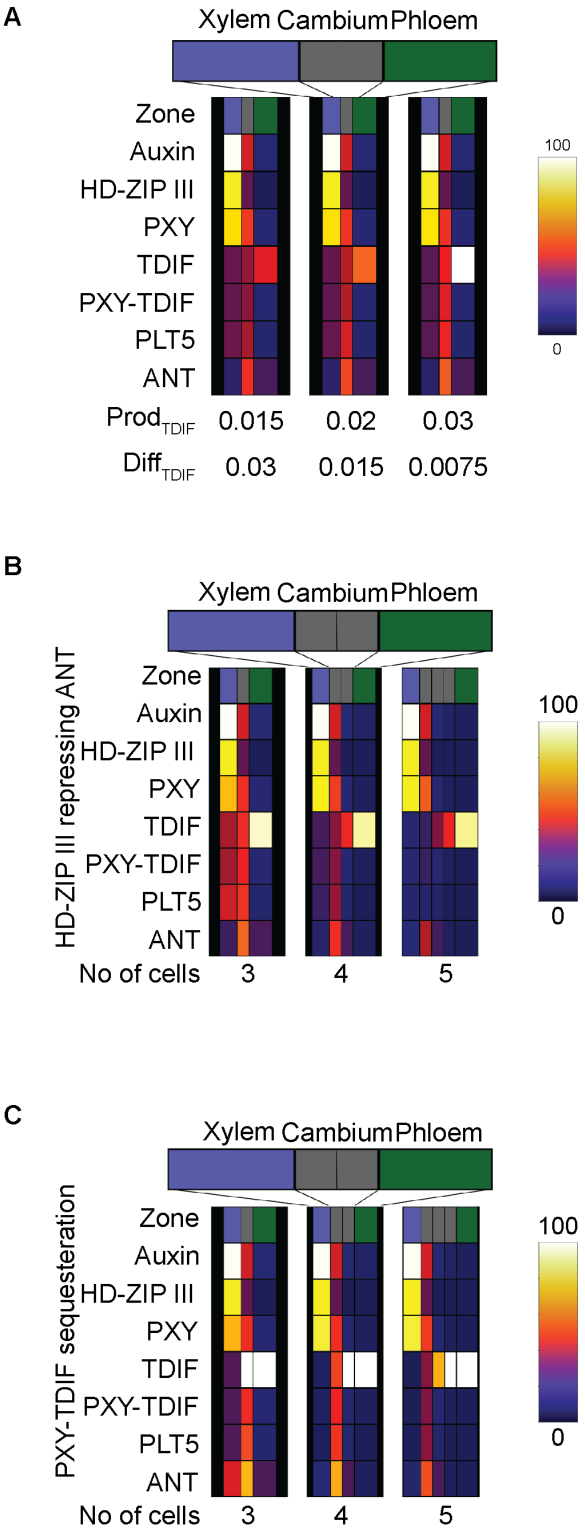
A
Xylem Cambium Phloem
Zone
Auxin
HD-ZIP III
PXY
TDIF
PXY-TDIF
PLT5
ANT
$Prod_{TDIF}$ 0.015 0.02 0.03
$Diff_{TDIF}$ 0.03 0.015 0.0075
100
0
B
HD-ZIP III repressing ANT
Xylem Cambium Phloem
Zone
Auxin
HD-ZIP III
PXY
TDIF
PXY-TDIF
PLT5
ANT
No of cells 3 4 5
100
0
C
PXY-TDIF sequesteration
Xylem Cambium Phloem
Zone
Auxin
HD-ZIP III
PXY
TDIF
PXY-TDIF
PLT5
ANT
No of cells 3 4 5
100
0

**Fig. S8. Robustness to variable TDIF gradients and cambial size.**
**(A)** robust cell fate decision making in a 3-cell tissue for variable production and diffusion rates of TDIF for strong HD-ZIP III repression settings. **(B)** and **(C)** robustness of cell fate decision making to variation in cambial cell number for the strong HD-ZIP III repression (A) and strong TDIF sequestration (C) regimes.

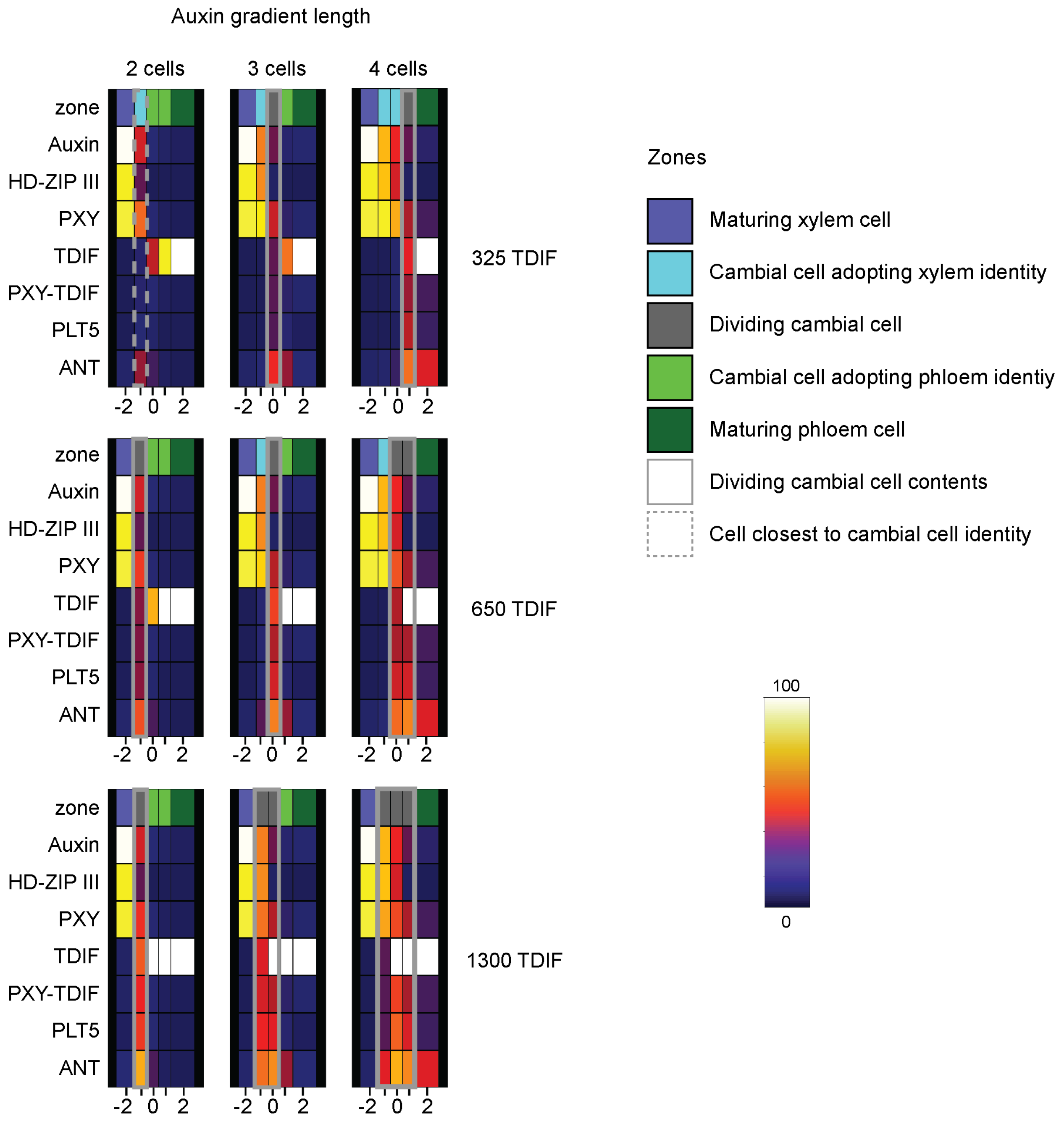
Auxin gradient length
2 cells
3 cells
4 cells
zone
Auxin
HD-ZIP III
PXY
TDIF
PXY-TDIF
PLT5
ANT
-2 0 2
325 TDIF
650 TDIF
1300 TDIF
Zones
Maturing xylem cell
Cambial cell adopting xylem identity
Dividing cambial cell
Cambial cell adopting phloem identiy
Maturing phloem cell
Dividing cambial cell contents
Cell closest to cambial cell identity
100
0

**Fig. S9. Flexibility to respond to variation in Auxin and TDIF gradients in 5 cell cambium**
Overview of cell fate decision making in a 5-cell vasculature exposed to variable auxin gradient lengths (left to right) and variable TDIF production levels in the phloem (top to bottom) under strong sequestration parameter settings. Gray bars show which cells will retain cambial identity (grey zone) as a result of expressing sufficient ANT+PLT. Light blue cells express sufficient HD-ZIP III to differentiate to xylem, while light green cells lack both high levels of ANT+PLT and HD-ZIP III and will thus differentiate to phloem. A strong auxin gradient (top right) pushes the cambial cell identity towards the phloem, allowing differentiation of new xylem cells. A strong TDIF signal pushes the cambial cell identity towards the xylem (bottom left) allowing phloem cells to differentiate. A combination of these two gradients allows for a larger total overlap (bottom right) that generates a larger cambium. When the overlap is too weak (top left) a meristem cannot be sustained. The dashed line indicates the highest current ANT+PLT level, where a smaller cambium (with increased PXY-TDIF overlap) could be maintained.

**table S1. Constant valued model parameters, their values and units.**

| Parameter name | Parameter value | Unit |
|---|---|---|
| $p_{HDZIPIII}$ | 0.02 | $[]s^{-1}$ |
| $d_{HDZIPIII}$ | 0.0002 | $s^{-1}$ |
| *pANT* | 0.002 | $[]s^{-1}$ |
| *dANT* | 0.00002 | $s^{-1}$ |
| *pPLT* | 0.002 | $[]s^{-1}$ |
| *dPLT* | 0.00002 | $s^{-1}$ |
| $ass_{PXYTDIF}$ | 0.02 | $[]^{-1}s^{-1}$ |
| $diss_{PXYTDIF}$ | 0.1 | $s^{-1}$ |
| $p_{PXY}$ | 0.02 | $[]s^{-1}$ |
| $d_{PXY}$ | 0.0012 | $s^{-1}$ |
| $d_{PXYTDIF}$ | 0.0012 | $s^{-1}$ |
| $d_{TDIF}$ | 0.0002 | $s^{-1}$ |

**table S2. Auxin dependent parameters, the range of values investigated, sampling interval used, values used for final model settings and units.**

| Parameter name | Parameter range | Interval | Final value | Unit |
|---|---|---|---|---|
| $Km_{Auxin,PXY}$ | 15-35 | 5 | 25 | [] |
| $Km_{Auxin,HDZIP}$ | 30-70 | 10 | 55 | [] |
| $Km_{Auxin,ANT}$ | 15-35 | 5 | 20 | [] |
| $Km_{HDZIP,ANT}$ | 30-50 | 5 | 50 | [] |
| $max_{Aux,ANT}$ | 0.1-0.4 | 0.05 | 0.2 | Dimensionless |
| $max_{PXY,ANT}$ | 0.6-0.9 | 0.05 | 0.8 | Dimensionless |
| $rep_{HDZIP,Aux}$ | 0.4-1. | 0.1 | 1.0 | Dimensionless |
| $rep_{HDZIP,PXY}$ | 0.4-1. | 0.1 | 1.0 | Dimensionless |

**table S3. PXY-TDIF dependent parameters, the range of values investigated, sampling interval used, values used for final model settings and units.**

| Parameter name | Parameter range | Interval | Final value | Unit |
|---|---|---|---|---|
| $Km_{PXYTDIF,ANT}$ | 15-35 | 5 | 20 | [] |
| $Km_{PXYTDIF,PLT}$ | 20-40 | 5 | 40 | [] |
| $Km_{PLT,PXY}$ | 30-50 | 5 | 50 | [] |
| $fac_{PLT,PXY}$ | 0.3 | 0 | 0.3 | [] |

**table S4. Diffusion parameters**

| Parameter name | Parameter values | Unit |
|---|---|---|
| $D_{TDIF}$ | 0.015 | μm $s^{-1}$ |
| $D_{ANT}$ | 0.00002 | μm $s^{-1}$ |
| $D_{PLT}$ | 0.00002 | μm $s^{-1}$ |

**table S5. Alternative parameter regimes**

| Parameter name | Strong HD-ZIP III repression value | Strong sequestering value | Unit |
|---|---|---|---|
| $p_{TDIF}$ | 0.03 | 0.13 | $[]s^{-1}$ |
| $rep_{HDZIP,Aux}$ | 1.0 | 1.0 | Dimensionless |
| $rep_{HDZIP,PXY}$ | 1.0 | 0.0 | Dimensionless |
| Effective TDIF degradation rate when bound to PXY | 0 | 0.0012 | $s^{-1}$ |
| $diss_{PXYTDIF}$ | 0.02 | 0.02 | $s^{-1}$ |

**table S6. Auxin and TDIF gradient values**

| | | | |
|---|---|---|---|
| Auxin gradient | | | |
| Gradient strength | Weak | Intermediate | Strong |
| $drop_{xylem}$ | 60 | 30 | 20 |
| $mod_{cambium}$ | 1.25 | 0.75 | 0.4 |
| TDIF gradient | | | |
| Gradient strength | Weak | Intermediate | strong |
| $p_{TDIF}$ | 0.065 | 0.13 | 0.26 |

**Data S1**

List of Primers, constructs, seeds
